# SLIT2-ROBO signaling in tumor-associated microglia/macrophages drives glioblastoma immunosuppression and vascular dysmorphia

**DOI:** 10.1101/2021.04.12.438457

**Authors:** Luiz Henrique Geraldo, Yunling Xu, Laurent Jacob, Laurence Pibouin Fragner, Rohit Rao, Nawal Maissa, Maïté Verreault, Nolwenn Lemaire, Camille Knosp, Corinne Lesaffre, Thomas Daubon, Joost Dejaegher, Lien Solie, Justine Rudewicz, Thomas Viel, Bertrand Tavitian, Steven De Vleeschouwer, Marc Sanson, Andreas Bikfalvi, Ahmed Idbaih, Q. Richard Lu, Flavia Regina Souza Lima, Jean-Leon Thomas, Anne Eichmann, Thomas Mathivet

## Abstract

SLIT2 is a secreted polypeptide that guides migration of cells expressing ROBO1&2 receptors. Herein, we investigated SLIT2/ROBO signaling effects in gliomas. In patients with glioblastoma (GBM), *SLIT2* expression increased with malignant progression and correlated with poor survival and immunosuppression. Knockdown of *SLIT2* in mouse glioma cells and patient derived GBM xenografts reduced tumor growth and synergized with immunotherapy to prolong survival. Tumor cell *SLIT2* knockdown inhibited macrophage invasion and promoted a cytotoxic gene expression profile, which improved tumor vessel function and enhanced efficacy of chemotherapy and immunotherapy. Mechanistically, SLIT2 promoted microglia/macrophage chemotaxis and tumor-supportive polarization via ROBO1&2-mediated PI3Kγ activation. Macrophage *Robo1&2* deletion and systemic SLIT2 trap delivery mimicked *SLIT2* knockdown effects on tumor growth and the tumor microenvironment (TME), revealing SLIT2 signaling through macrophage ROBOs as a novel regulator of the GBM microenvironment and a potential immunotherapeutic target for brain tumors.

## Introduction

Malignant gliomas are the most common primary brain tumors (1, 2). Among those, Glioblastoma (GBM, WHO grade IV glioma) is the most frequent and aggressive tumor that accounts for more than 50% of gliomas, with poor patient prognosis (3). GBMs are molecularly heterogeneous and invasive, angiogenic and proliferative tumors that are largely resistant to current therapies (4).

Tumor-associated Microglia and Macrophages (TAMs) are the most abundant cells in the GBM microenvironment, composing up to 25% of the tumor mass (5–7). TAMs are key drivers of GBM immunosuppression and pathological angiogenesis (7). TAMs inhibit T cell responses in the GBM microenvironment by favoring regulatory T cells and suppressing anti-tumor T cell responses (8–11), thereby limiting the efficacy of currently available T cell-oriented immunotherapies in GBM (4, 12–14). TAM-derived signaling also contributes to vascular dysmorphia, and drives blood vessel dilation and leakiness in the GBM microenvironment (15, 16). Non-uniform oxygen delivery via dysmorphic and leaky tumor vessels leads to hypoxia, which upregulates angiogenic factors that induce more dysfunctional vessels, thereby preventing the delivery of cytotoxic agents to kill tumor cells (4, 17). The mechanisms by which TAMs promote vessel dysmorphia and immune evasion are as yet incompletely understood, and the means to prevent them are not available (7, 18, 19).

SLITs are evolutionary conserved secreted polypeptides that bind to transmembrane Roundabout (ROBO) receptors (20, 21). In mammals, three SLIT ligands (SLIT1-3) signal via two ROBO receptors, ROBO1 and 2 (22). SLIT ligands bind via the second leucine-rich repeat region (D2) to the Ig1 domain of ROBO1&2 (23), while mammalian ROBO3 and ROBO4 lack the SLIT binding residues and do not bind SLITs (24, 25). SLIT binding triggers recruitment of adaptor proteins to the ROBO cytoplasmic domain that modulate the cytoskeleton, in turn regulating cell migration, adhesion and proliferation (22, 26, 27).

SLIT-ROBO signaling was discovered in the developing nervous system as a guidance cue for axonal growth cones that regulates pathfinding of commissural axons and motor coordination between the left and right side of the body (20, 21). It is now known that SLIT-ROBO signaling controls several additional biological processes, including angiogenesis and immune cell migration.

In endothelial cells, SLIT2 activation of ROBO1&2 signaling promotes retinal and bone angiogenesis by driving tip cell migration and polarization (28–31). In the immune system, SLITs have been described as chemo-attractive for neutrophils (32) and chemorepellent for lymphocytes and dendritic cells (33–36). In macrophages, SLIT-ROBO signaling prevented macropinocytosis and cytotoxic polarization (37).

In tumor contexts, SLIT2 exerts a pro-angiogenic role (38–40), and has been reported to enhance tumor cell aggressiveness and migration (41–45), metastatic spread (40, 46) and therapy resistance (47), particularly in colorectal cancer, pancreatic cancer and osteosarcoma. Nevertheless, other studies reported a tumor suppressive role for SLIT2-ROBO signaling in in lung and breast cancers (48–50). In the context of GBM, some studies suggested that SLIT2 signaling could inhibit tumor growth (51–53), while in others SLIT-ROBO signaling correlated with more aggressive GBM behavior (54, 55). Given the various and context dependent effects of SLIT-ROBO signaling in cancer, it remained unclear if this pathway could be used therapeutically to prevent cancer growth.

We showed here that high SLIT2 expression in GBM patients and in mouse models induced TAM accumulation and vascular dysmorphia, and that *SLIT2* knockdown in glioma cells and systemic SLIT2 inhibition with a ligand trap normalized the TME by preventing TAM tumor-supportive polarization and angiogenic gene expression. As a result, anti-tumor immune responses and tumor perfusion were enhanced, and efficacy of temozolomide(TMZ)-based chemotherapy and checkpoint inhibitor-based immunotherapy were increased. Inducible genetic deletion of *Robo1&2* in macrophages was sufficient to normalize the TME and enhanced response to immunotherapy, revealing a novel macrophage-based immunotherapy approach for GBM.

## Results

### *SLIT2* expression correlated with poor glioma patient prognosis

The Cancer Genome Atlas (TCGA) RNA sequencing data analysis showed that high *SLIT2* expression was significantly associated with decreased GBM patient survival (Figure 1A, O.S., 9.86 months for high expression, 14.69 months for low expression, and 16.79 months for medium expression, log-rank test), whereas higher expression of the other *SLIT* family members and *ROBO* receptors was not (Supplemental Figure 1A-D). Analysis of TCGA Agilent-4502A microarray confirmed that high *SLIT2* expression correlated with poor survival (Supplemental Figure 1E, O.S., 12.9 months for high expression and 15.1 months for low expression). Analysis of a primary glioma patient cohort (129 patients, 84 Low Grade Gliomas and 45 GBMs) also demonstrated correlation between high *SLIT2* expression and worse prognosis in both low-grade gliomas (LGGs) and GBMs (Figure 1B, O.S., for LGG: 64.73 months for high expression and 209.10 months for low expression; and Supplemental Figure 1F, O.S., for GBM: 14.75 months for high expression and 16.25 months for low expression).

**Figure 1.**
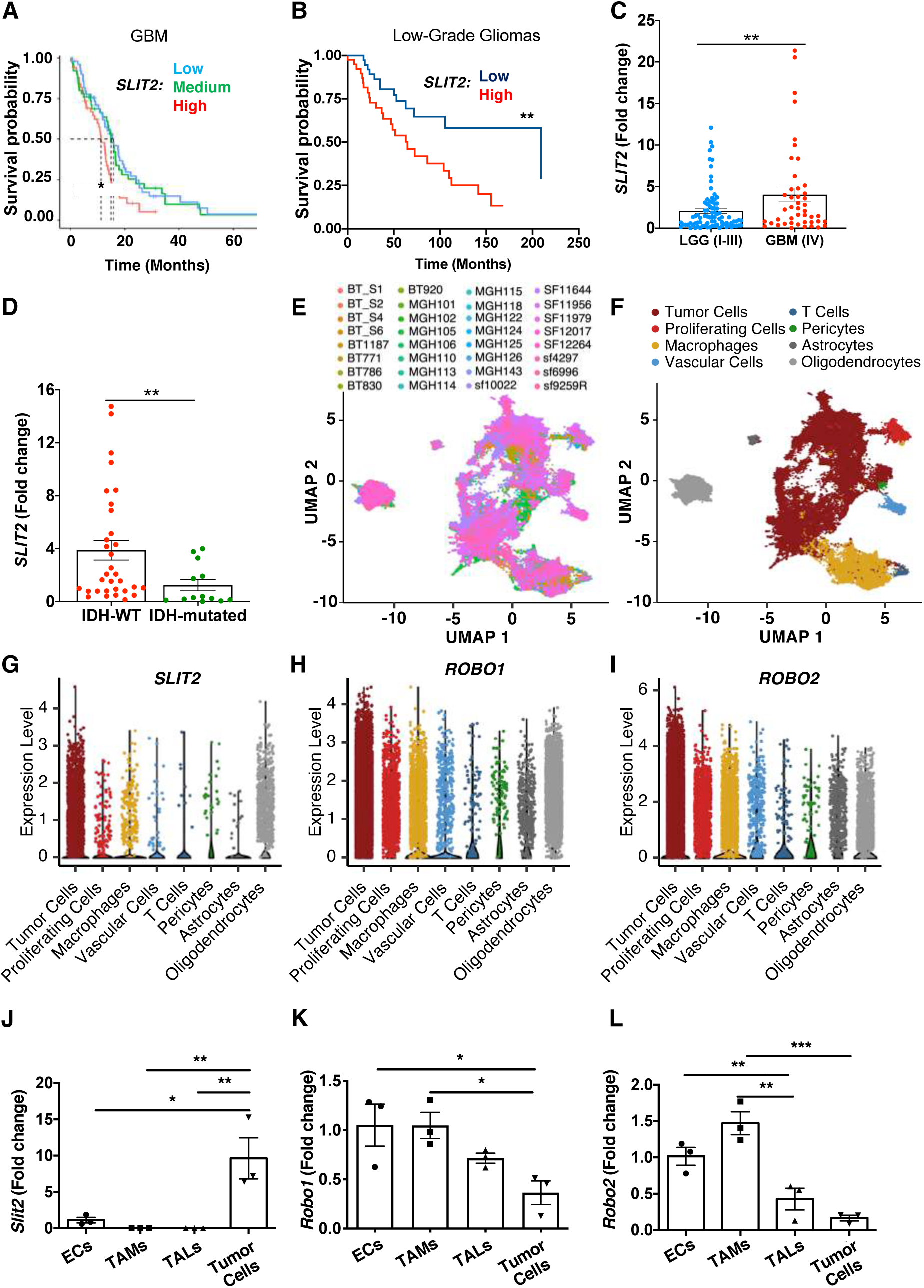
Slit2 expression correlates with glioma aggressiveness and poor patient prognosis. **A.** *In silico* analysis of TCGA glioblastoma RNAseq patient database (*n =* 51 high, 50 medium and 50 low Slit2 expressing patients; O.S., 9.86 months for high expression, 14.69 months for low expression, and 16.79 months for medium expression, log-rank test). **B.** Survival analysis of Low-Grade Gliomas (LGG, Grades I to III) patients grouped by their levels of *SLIT2* expression (*n =* 41 high and 41 low *SLIT2* expressing patients; O.S., 64.73 months for high expression and 209.13 months for low expression, log-rank test). **C.** *SLIT2* qPCR expression in glioma patient samples from (**B)** (GBM, *n =* 45; LGG, *n =* 84; Student’s t test). **D.** *SLIT2* qPCR expression in Grades III and IV glioma patient samples classified by their IDH-1/2 status (IDH-WT, *n =* 51; IDH-mutated, *n =* 34; Mann-Whitney U test). **E-F.** UMAP plots of single cell RNA sequencing (scRNAseq) of 32 GBM patients showing different samples (**E**) and clustering of the different cell types in the GBM microenvironment (**F**). **G-I.** Expression plots of *SLIT2* (**G**), *ROBO1* (**H**) and *ROBO2* (**I**) in scRNAseq data from (**E)**. **J-L.** qPCR analysis of *Slit2* (**J**), *Robo1* (**K**) and *Robo2* (**L**) expression in endothelial cells (ECs), tumor associated macrophages (TAMs), tumor associated T lymphocytes (TALs) and tumor cells FACS-sorted from late-stage CT-2A mice glioblastomas (*n =* 3 independent tumors, day 21 after implantation, One-Way ANOVA). All data are represented as mean ± s.e.m. * *P* < 0.05, ** *P* < 0.01, *** *P* < 0.001.

Further analysis of RNA sequencing data demonstrated higher *SLIT2* expression in the most aggressive and angiogenic mesenchymal GBM subtype (Verhaak et al., 2010) and lower expression in classical GBMs (Supplemental Figure 1G). High *SLIT2* expression was also associated with poor survival in patients with mesenchymal GBM in this cohort (Supplemental Figure 1H, O.S., 10.4 months for high expression and 17.9 months for low expression, log-rank test). Finally, qPCR analysis of patient samples also revealed higher expression levels of *SLIT2* in WHO grade IV GBM compared to WHO grade I, II and III glioma patients (Figure 1C), while expression of other *SLITs* and *ROBOs* was not changed between glioma grades (Supplemental Figure 1I-L). Expression levels of *SLIT1* and *SLIT3* were significantly lower compared to *SLIT2* in GBM patients (Supplemental Figure 1M).

Isocitrate dehydrogenase 1 and 2 (IDH-1/2) mutations are known prognostic factors in malignant gliomas. Patients with grade III gliomas and no IDH mutations (IDH-WT) have comparable prognosis to those of GBM patients, while patients with IDH mutations have better survival prognosis (56–58). We compared glioma patients classified by IDH-status, and observed increased *SLIT2* expression in patients with IDH-WT tumors in either grade III and IV gliomas (Figure 1D) or in all gliomas (Supplemental Figure 1N).

To determine the source of *SLIT2* in the GBM microenvironment, we analyzed single cell RNA sequencing (scRNAseq) data from human GBM patients (Figure 1E-F). *SLIT2* expression was highest in tumor cells and oligodendrocytes (Figure 1G), while *ROBO1* and *ROBO2* expression were highest in tumor cells but also detected in other cell types in the TME, particularly in TAMs (Figure 1H-I).

We next generated a mouse model of GBM by intra-cerebral inoculation of syngenic CT-2A mouse glioma tumor cells expressing green fluorescent protein (GFP) into adult C57BL/6 mice (16, 59). Expression of *Slit* ligands and their *Robo* receptors was tested 21 days after tumor cell inoculation by qPCR on FACS-sorted tumor cells (GFP^+^), endothelial cells (ECs, CD31^+^), TAMs (CD45^+^CD11b^+^CD3^-^), and Tumor-associated T Lymphocytes (TALs, CD45^+^CD11b^-^CD3^+^). The major source of *Slit2* were the tumor cells themselves (Figure 1J). By contrast, *Robo1* and *Robo2* receptors were mainly expressed by ECs and recruited TAMs and TALs (Figure 1K-L). *Slit1* and *Slit3* expression levels in mouse tumor cells were much lower when compared to *Slit2* (Supplemental Figure 1O). These data suggested that interactions between tumor cell-derived SLIT2 and stromal cells expressing ROBOs could affect GBM growth.

### *Slit2* silencing slowed GBM growth and increased TMZ sensitivity

To determine if tumor cell-derived Slit2 affected GBM growth, we infected CT-2A and GL261 glioma cells with lentivirus encoding GFP-tagged control scrambled shRNA (shCTRL) or *Slit2* targeting shRNA (shSlit2) alone or combined with an shRNA-resistant human SLIT2 construct (shSlit2 + hSLIT2). *Slit2* knockdown significantly decreased Slit2 protein and mRNA expression, while shSlit2 + hSLIT2 cells expressed more Slit2 than controls (Figure 2A-B and Supplemental Figure 2A-F). Expression of other Slits or Robo1 and 2 was not altered (data not shown). *In vitro* growth rates of shCTRL and shSlit2 CT2A and GL261 knockdown cells were similar (Supplemental Figure 2G-H). Slit2 did not induce tumor cell chemotaxis in a transwell chamber assay (Supplemental Figure 2I-J). Nevertheless, migration of shSlit2 cells towards a serum gradient in the lower chamber was reduced (Supplemental Figure 2K-L).

**Figure 2.**
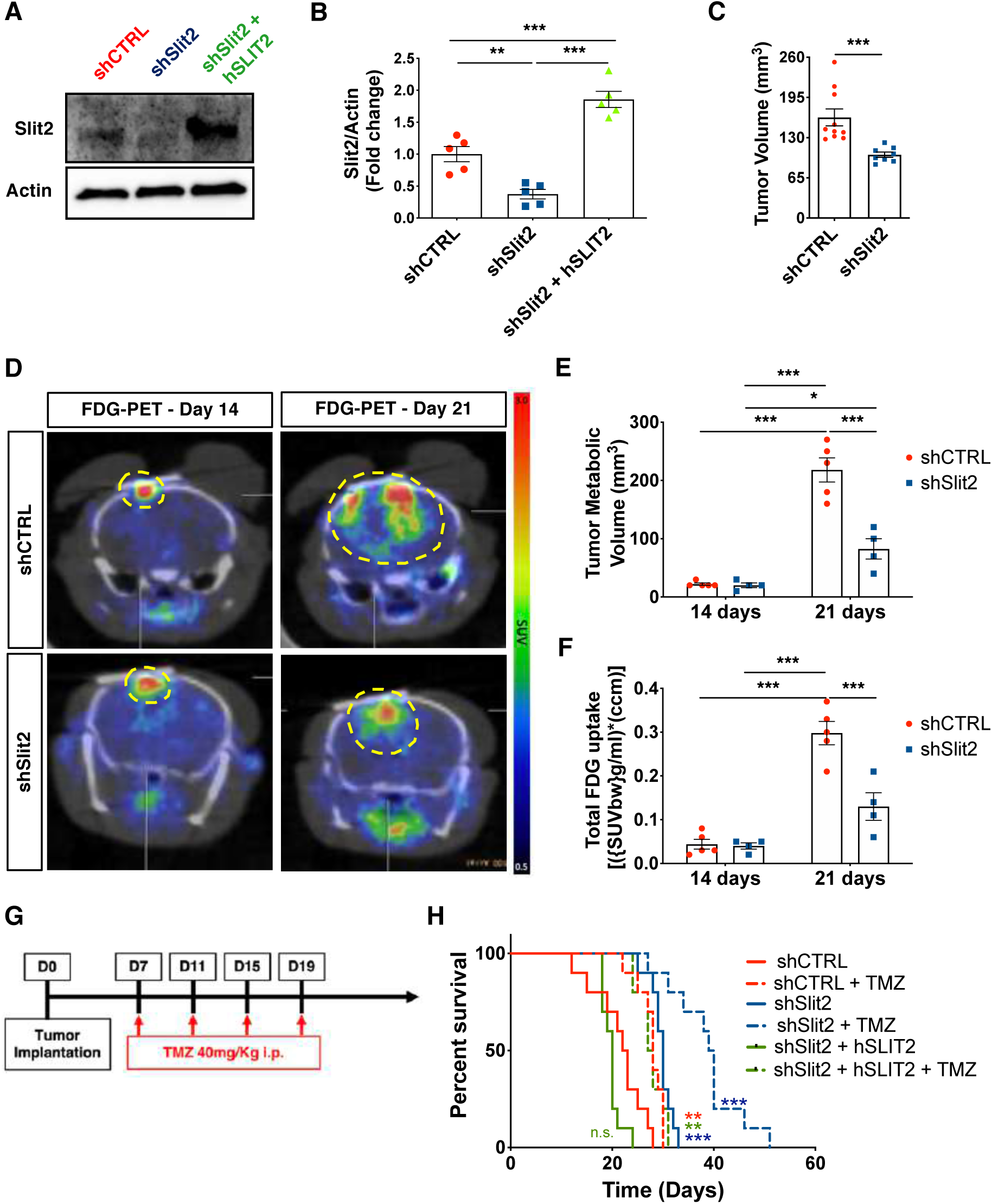
Slit2 promotes Glioblastoma growth and resistance to TMZ. **A-B.** Western blot analysis (**A**) and quantification (**B**) of Slit2 expression in shCTRL, shSlit2 and shSlit2+hSLIT2 CT-2A cells (*n =* 5, One-Way ANOVA). **C.** Tumor volume quantification at 21 days (*n =* 10 for shCTRL and *n =* 8 for shSlit2, Student’s t-test). **D.** FDG-PET imaging over CT-2A shCTRL and shSlit2 glioma growth (*n =* 5 shCTRL and *n =* 4 shSlit2). **E-F.** Quantification of tumor metabolic volume (**E**) and total tumor glucose uptake (**F**) from **(D)** (*n =* 5 for shCTRL and *n =* 4 for shSlit2, One-Way ANOVA). **G.** Survival trial design: 8-week-old mice were engrafted with CT-2A shCTRL, shSlit2 or shSlit2+hSLIT2 spheroids and randomly assigned to vehicle or TMZ treatment (40 mg/kg on days 7, 11, 15 and 19 after tumor implantation). **H.** Survival curves of the mice in **(G)** (*n =* 10 mice per group, O.S.= 22.5 days for shCTRL, 28 days for shCTRL + TMZ, 30 days for shSlit2, 39.5 days for shSlit2 + TMZ, 20 days for shSlit2+hSLIT2 and 27 days for shSlit2+hSLIT2 + TMZ; Multiple comparisons log-rank test). Data are presented as mean ± s.e.m. * *P* < 0.05, ** *P* < 0.01, *** *P* < 0.001

Individual 250-μm diameter tumor cell spheroids were implanted through cranial windows into Tomato-fluorescent reporter mice (ROSA^mT/mG^) and followed longitudinally. Compared to shCTRL, Slit2 knockdown tumors exhibited reduced volumes after 21 days (Figure 2C). FDG-PET imaging showed that tumor metabolic volume and FDG total uptake were similar between shSlit2 and shCTRL at 14 days, but reduced in shSlit2 tumors at 21 days (Figure 2D-F), demonstrating that *Slit2* knockdown delayed tumor growth *in vivo*.

We investigated if *Slit2* knockdown affected survival in combination with low-dose chemotherapy with the DNA alkylating agent TMZ, a classical treatment for GBM (Figure 2G). Compared to shCTRL, *Slit2* knockdown increased overall survival of tumor-bearing mice, while Slit2 overexpression tended to decrease survival (Figure 2H, O.S., 22.5 days for shCTRL, 30 days for shSlit2 and 20 days shSlit2 + hSLIT2). TMZ treatment further increased overall survival of shSlit2 glioma-bearing mice (Figure 2H, O.S., 28 days TMZ for shCTRL+ TMZ, 39 days for shSlit2 + TMZ and 27.5 days for shSlit2 + hSLIT2 + TMZ). shSlit2 did not affect TMZ sensitivity of tumor cells in vitro (Supplemental Figure 2M), but significantly increased TMZ-induced pH2AX^+^ double strand DNA breaks in tumors *in vivo* (Supplemental Figure 2N-O), suggesting that changes in the TME might contribute to enhanced TMZ sensitivity *in vivo*.

### *SLIT2* silencing slowed GBM growth and invasiveness in a Patient-derived Xenograft model (PDX)

To determine whether SLIT2 had similar effects on human GBM tumors, we used N15-0460 patient-derived GBM cells that were established from a biopsy and grown as tumor spheres. We infected these cells with lentivirus encoding a luciferase reporter and GFP-tagged control scrambled shRNA (shCTRL) or *SLIT2* targeting shRNA (shSLIT2). *SLIT2* knockdown significantly decreased SLIT2 protein and mRNA expression, without altering expression of other *SLIT*s or *ROBO1* and *2* (Supplemental Figure 3A-G). *In vitro* growth rates of shCTRL and shSLIT2 cells and sensitivity to TMZ were similar (Supplemental Figure 3H-I). SLIT2 did not induce tumor cell chemotaxis in a transwell chamber assay, but migration of shSLIT2 cells towards a serum gradient in the lower chamber was reduced (Supplemental Figure 3J-K). Next, we analyzed sphere formation, and observed that shCTRL and shSLIT2 cells formed similar numbers of spheres after 48hs in culture, but the size of shSLIT2 spheres was reduced when compared to shCTRL (Supplemental Figure 3L-M). Analysis of tumor sphere invasion in fibrin gels showed that shSLIT2 decreased spheroid invasion after 24 and 48 hours in culture when compared to shCTRL (Supplemental Figure 3N-O).

To determine the effect of shSLIT2 on human GBM growth, we implanted shCTRL and shSLIT2 N15-0460 cells in Hsd:Athymic Nude-Foxn1nu mice and followed tumor growth by bioluminescence analysis every 2 weeks after tumor implantation. 170 days after tumor implantation, 80% of the mice injected with shCTRL cells developed tumors, while only 20% of shSLIT2-injected mice had tumors (Supplemental Figure 4A). Analysis of the bioluminescence curves of shCTRL and shSLIT2 tumors demonstrated that more mice developed tumors in the shCTRL group, and that the shCTRL tumors were bigger than shSLIT2 tumors (Supplemental Figure 4B-C). Histological analysis of GFP^+^ tumor cells on vibratome sections 170 days after tumor implantation showed that shCTRL cells either developed tumor masses or spread throughout the entire brain, while shSLIT2 cells remained restrained to the injection site or migrated through the corpus callosum, but did not form tumor masses (Supplemental Figure 4D-E). *SLIT2* shRNA also reduced the expression of SOX2 and PML involved in GBM tumor cell malignancy (55, 60, 61) (Supplemental Figure 4F-H).

### *Slit2* knockdown improved tumor vessel function

To determine if tumor-secreted SLIT2 affected the GBM microenvironment, we used 2-photon *in vivo* imaging of red fluorescent ROSA^mT/mG^ mice. We observed that blood vessels in shCTRL CT2A and GL261 tumors became abnormally enlarged and lost branching points between day 14 and day 21, while shSlit2 tumor vessels dilated less and remained more ramified (Figure 3A-C, Supplemental Figure 5A-D). Conversely, SLIT2 overexpressing tumor vessels dilated and lost branchpoints earlier, at day 18 after injection (Figure 3D-F, Supplemental Figure 5E), just prior to their death at 20 days post tumor implantation.

**Figure 3.**
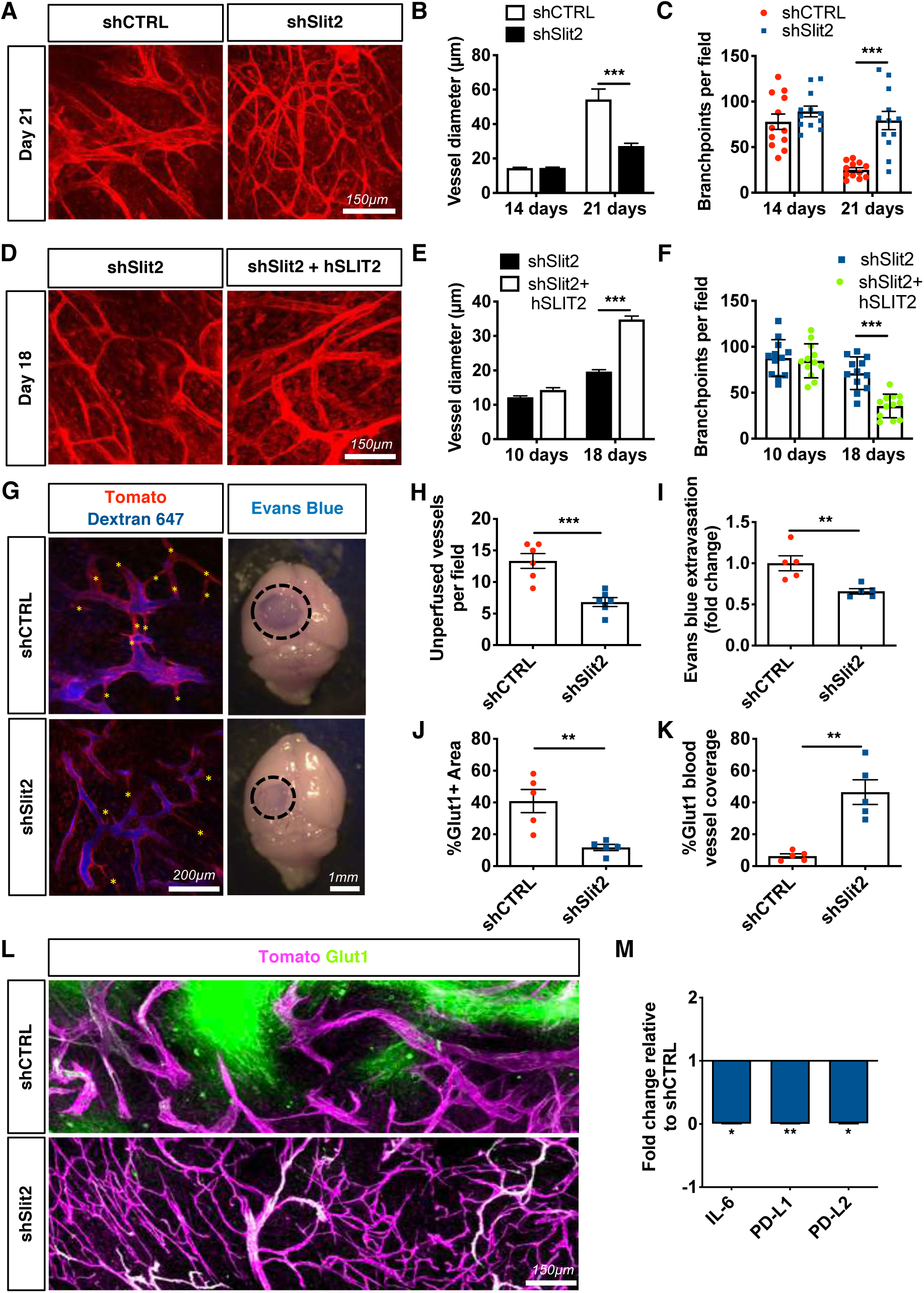
Slit2 promotes blood vessel dysmorphia in GBM. **A.** *In vivo* two-photon images of ROSA^mTmG^ mice bearing day 21 CT-2A shCTRL or shSlit2 tumors. **B-C.** Quantification of vessel diameter (**B**) and branchpoints (**C**) (*n =* 8 mice per group, One-way ANOVA). **D**. *In vivo* two-photon images of ROSA^mTmG^ mice bearing day 18 CT-2A shSlit2 or shSlit2+hSLIT2 tumors. **E-F**. Quantification of vessel diameter (**E**) and branchpoints (**F**) (*n =* 7 mice per group, One-way ANOVA). **G-I**. Left panels: Two-photon *in vivo* imaging following intravenous injection of Alexa Fluor 647 conjugated Dextran highlighting unperfused blood vessel segments in the tumor core (asterisks) of day 21 CT-2A shCTRL and shSlit2 tumors. Right panels: representative pictures of whole brains of day 21 shCTRL or shSlit2 CT-2A tumors following Evans blue injection. **H.** Quantification of unperfused blood vessel segments in the tumor mass presented in **(G)** (*n =* 5 mice per group, Mann-Whitney U test). **I.** Quantification of Evans Blue extravasation in **(G)** (*n =* 5 mice per group, Mann-Whitney U test). **J-L.** quantifications of Glut1+ hypoxic areas in the tumor (**J**) and Glut1 blood vessel coverage **(K)** from immunohistochemistry on sections (**L**) (*n =* 5 mice per group, Mann-Whitney U test). **M.** qPCR analyses from FACS-sorted endothelial cells (*n =* 3 tumors/group, Mann-Whitney U test). Data are presented as mean ± s.e.m. * *P* < 0.05, ** *P* < 0.01, *** *P* < 0.001

Functionally*, in vivo* imaging after intravenous Alexa Fluor 647-labeled dextran injection revealed significantly improved perfusion in shSlit2 CT2A tumor vessels when compared to shCTRL tumors (Figure 3G-H). Quantification of Evan’s blue extravasation showed reduced vascular leakage in shSlit2 tumors compared to shCTRL (Figure 3I). Along with improved vascular function in shSlit2 knockdown tumors, glucose transporter 1 immunostaining (Glut1)-positive hypoxic areas within the tumor mass were reduced, and Glut1 coverage of blood vessels was increased in shSlit2 knockdown tumors compared to shCTRL, indicating partially improved blood-brain barrier function (Figure 3J-L). QPCR analysis of sorted tumor endothelial cells (CD45^-^CD31^+^) showed downregulation of immunosuppressive IL-6, PD-L1 and PD-L2 in Slit2 shRNA transfected tumors compared to CTRL tumors (Figure 3M).

### *Slit2* silencing reduced myeloid immunosuppression

*In vivo* imaging also revealed that immune cell infiltration was increased in SLIT2 overexpressing tumors, and decreased in Slit2 silenced tumors when compared to CTRL tumors (Supplemental Figure 6A-C). Immunofluorescence analysis of tumor sections showed a decrease in the numbers of F4/80^+^ myeloid cells in day 21 shSlit2 tumors compared to day 21 shCTRL or day 18 SLIT2-overexpressing tumors (Figure 4A-B, Supplemental Figure 6D-E). Activated MHC-II^+^ antigen-presenting cells (APCs) were increased in shSlit2 tumors, and MRC1(CD206)^+^ tumor-supportive infiltrating immune cells were decreased (Figure 4A-B, Supplemental Figure 6D-E). FACS sorted CD45^+^CD11b^+^F4/80^+^Ly6G^-^ TAMs accounted for about 12% of the total cells in shCTRL tumors, but only 6% in the *Slit2* knockdown tumors (Figure 4C, Supplemental Figure 6F). Half of the TAMs in shSlit2 CT2A tumors had a cytotoxic activation profile and expressed MHC-II and CD11c, while <20% of TAMs in the shCTRL condition expressed MHCII and CD11c and >80% expressed the tumor supportive marker MRC1 (Figure 4D). shSlit2 tumors also showed increased infiltration of Dendritic Cells (CD45^+^CD11b^-^CD11c^+^MHC-II^+^F4/80^-^) and neutrophils (CD45^+^CD11b^+^Ly6G^+^) that were much less abundant when compared to TAMs (Supplemental Figure 6G-H).

**Figure 4.**
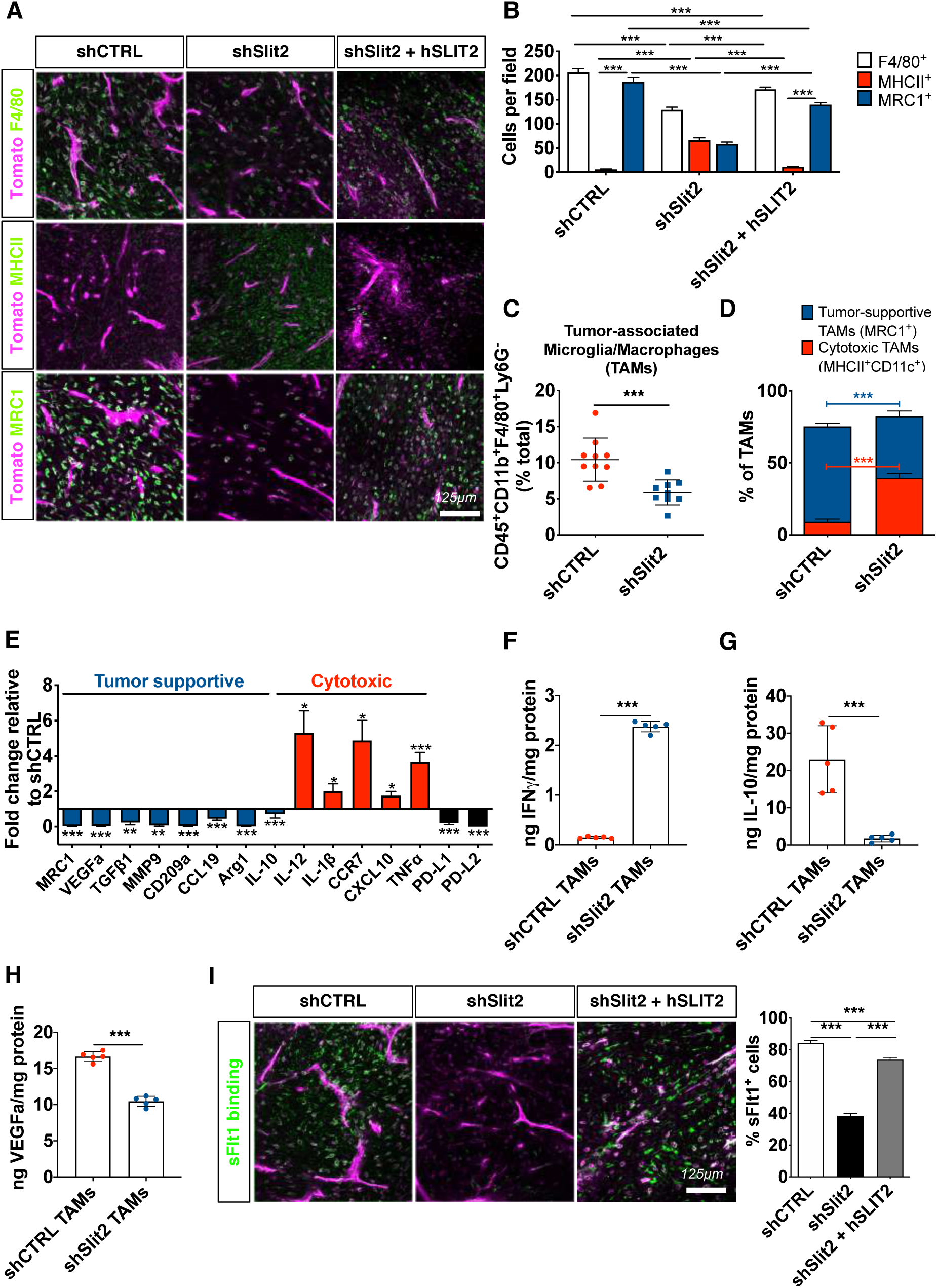
Slit2 promotes TAM recruitment and polarization in mouse gliomas. **A.** Immunohistochemistry on sections of late-stage CT-2A shCTRL, shSlit2 or shSlit2+hSLIT2 tumors for F4/80, MHC-II and MRC1^+^ cells (green). **B**. Quantifications of (**A)** (*n =* 7 mice per group, 5 fields per tumor, Two-Way ANOVA). **C-D.** FACS analysis of 21 days CT-2A shCTRL and shSlit2 for quantification of TAMs (*n =* 10 tumors/group; Student’s t-test and Two-way ANOVA). **E.** qPCR analysis from FACS- sorted TAMs (*n =* 6 tumors/group, Mann-Whitney U test). **F-H.** ELISA from protein samples extracted from FACS-sorted TAMs from shCTRL and shSlit2 tumors to quantify IFNγ (**F**), IL-10 (**G**) and VEGFa (**H**) (*n =* 5 tumors/group, Mann-Whitney U test). **I.** Representative images and quantification of soluble-Flt1 binding to sections of day 21 CT-2A shCTRL, shSlit2 and day 18 shSlit2+hSLIT2 tumors (*n =* 7 mice per group, 5 fields per tumor, One-Way ANOVA). * *P* < 0.05, ** *P* < 0.01, *** *P* < 0.001

Molecularly, when compared to FACS-sorted shCTRL, TAMs from shSlit2 tumors exhibited decreased expression of tumor supportive genes *Mrc1*, *Vegfa*, *Tgfβ1*, *Mmp9*, *Cd209a*, *Ccl19*, *Arg1* and *Il10*, increased expression of cytotoxic genes *Il-12*, *Il-1b*, *Ccr7*, *Cxcl10* and *Tnfα*, and reduced expression of *Pd-l1* and *Pd-l2* inhibitors of T cell activation (Figure 4E). ELISA analysis showed increased IFNγ and confirmed reduced IL-10 and VEGFa protein levels in TAMs sorted from shSlit2 tumors when compared to controls (Figure 4F-H). In line with reduced VEGFa expression, *in vivo* binding of soluble VEGFR1 (sFlt1) showed that only about 40% of stromal cells in shSlit2 tumors bound sFlt1, while >80% of CTRL and SLIT2 overexpressing cells bound Flt1 (Figure 4I).

### SLIT2 inhibition increased T cell infiltration and improved checkpoint inhibitor treatment

In contrast to the decreased number of TAMs in shSlit2 tumors, the total number of TALs was increased 3-fold (Figure 5A, Supplemental Figure 7A-D), with an increase in both CD4^+^ and CD8^+^ T lymphocytes within the tumor mass when compared to controls (Figure 5B-C and Supplemental Figure 7E-I). Furthermore, the CD4^+^ TALs in shSlit2 tumors showed increased expression of Th1 response related genes (*Ifnγ, Cxcl11* and *Il-2*) and of IL-17a, but decreased expression of Th2 response related genes (*Il-10* and *Cxcl10*) and PD-1 and CTLA4 (Figure 5D). CD8^+^ TALs in shSlit2 tumors also showed increased expression of activation markers (IFN*γ* and GZMB), and reduced expression of genes related to CD8 T cell exhaustion (Tim3 and Lag3) (62) (Figure 5E). In tumor sections, we observed more infiltrating GZMB^+^ CD8^+^ activated anti-tumor T cells in shSlit2 compared to shCTRL tumors (Figure 5F-G). ELISA analysis of these sorted CD8^+^ TALs also showed increased IFNγ (Figure 5H) and reduced IL-10 and VEGFa protein levels (Figure 5I-J) in cells sorted from shSlit2 tumors.

**Figure 5.**
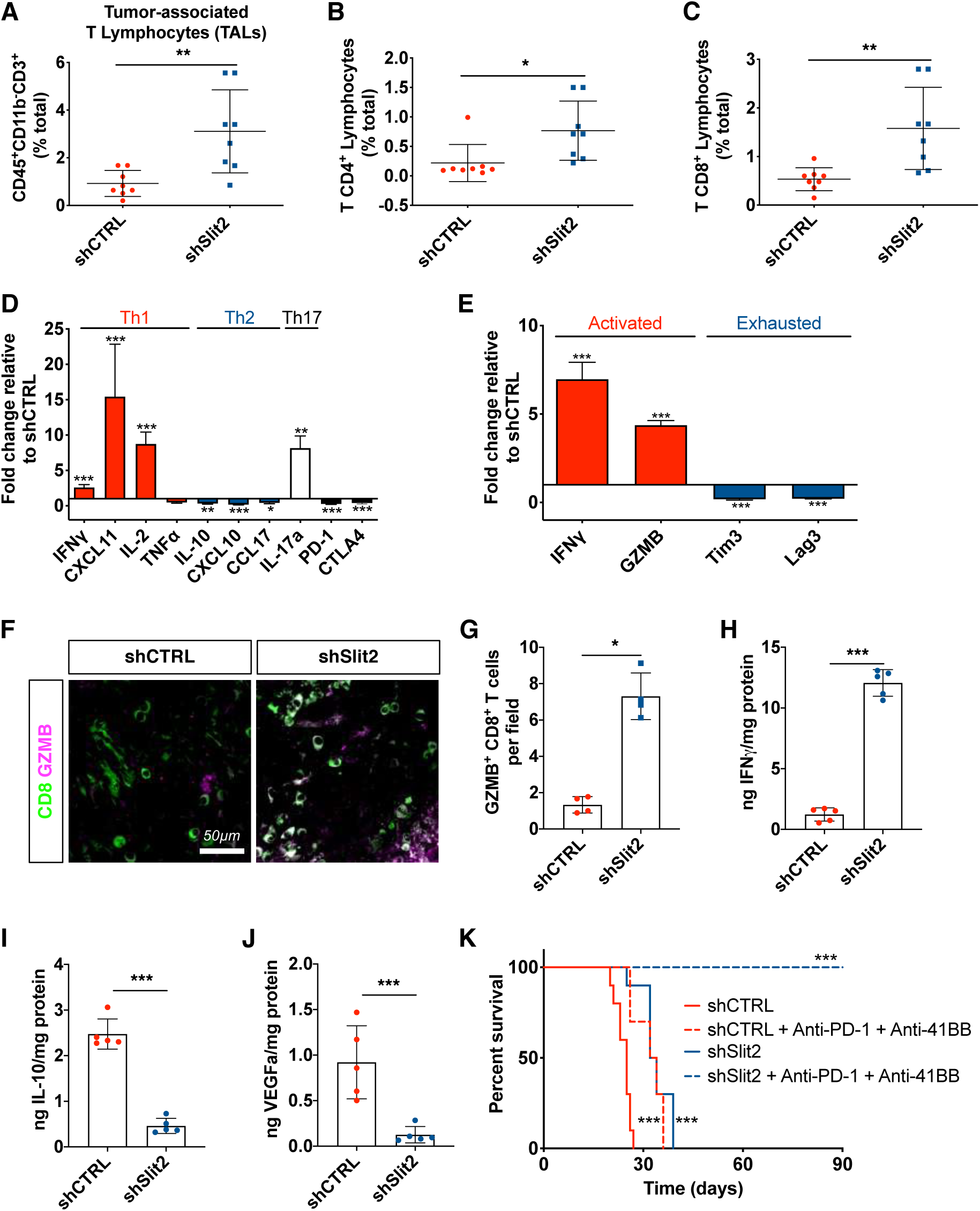
Slit2 inhibits T lymphocyte responses in the glioma microenvironment. **A-C**. T lymphocyte FACS analysis of day 21 CT-2A shCTRL and shSlit2 tumors for total CD3^+^ TALs (**A**), CD4^+^ TALs (**B**) and CD8^+^ TALs (**C**) (*n =* 8 tumors/group; Student’s t-test). **D.** qPCR analyses from FACS-sorted CD4^+^ T lymphocytes (*n =* 10 tumors/group, Mann-Whitney U test). **E.** qPCR analyses from FACS-sorted CD8^+^ T lymphocytes (*n =* 6 tumors/group, Mann-Whitney U test). **F.** Representative images of CD8 and GZMB staining on sections of day 21 CT-2A shCTRL and shSlit2 tumors. **G.** Quantification of **(F)** (*n =* 4 mice per group, 5 fields per tumor, Mann-Whitney U test). **H-J.** ELISA from protein samples extracted from FACS-sorted CD8^+^ TALs from shCTRL and shSlit2 tumors to quantify IFNγ (**H**), IL-10 (**I**) and VEGFa (**J**) (*n =* 5 tumors/group, Mann-Whitney U test). **K.** 8-week-old mice were engrafted with CT-2A shCTRL or shSlit2 and randomly assigned to vehicle or Anti-PD-1 + Anti-4-1BB treatment (200ug each on days 7, 9, 11 and 13 after tumor implantation) (*n =* 10/11 mice per group, O.S.= 25 days for shCTRL, 33 days for shCTRL + Anti-PD-1 + Anti-4-1BB, 33 days for shSlit2 and Undetermined for shSlit2 + Anti-PD-1 + Anti-4-1BB; Multiple comparisons log-rank test). Data are presented as mean ± s.e.m. * *P* < 0.05, ** *P* < 0.01, *** *P* < 0.001

Given this shift towards a less immunosuppressive GBM microenvironment, we hypothesized that shSlit2 tumors would be more sensitive to treatment with immune checkpoint inhibitors using anti-PD-1 and anti-4-1BB antibodies (11, 59). We treated mice with 200ug of each antibody at D7, D9, D11 and D13 after tumor implantation. Combining immune checkpoint inhibitor therapy with *Slit2* silencing led to powerful anti-tumor responses, with 100% of the mice alive at 90 days after implantation (Figure 5K, O.S.= 25 days for shCTRL, 33 days for shCTRL + Anti-PD-1 + Anti-4-1BB, 33 days for shSlit2 and Undetermined for shSlit2 + Anti-PD-1 + Anti-4-1BB).

The changes in the immune cell microenvironment that we observed in the murine GBM models are also likely to occur in GBM patients, as shown by positive correlation between *SLIT2* and *MRC1* and *VEGFA* mRNA expression in patient samples from our GBM patient cohort and TCGA database cohorts (Supplemental Figure 8A-C). *SLIT2* expression also correlated with genes related to tumor-supportive macrophages (*CCL19*, *CD209*, *MMP9* and *PD-L2*), inhibition of anti-tumor T cell responses (*PD-1, CTLA4, CCL17, CXCL11*, *LAG3* and *TIM3*) and *IL-6* for example (Supplemental Figure 8D-O).

### SLIT2 promoted microglia and macrophage migration and polarization via ROBO1&2

To determine how SLIT2 affected myeloid cells, we tested microglia and macrophage migration in Transwell chambers. Slit2 in the bottom chamber induced chemotaxis of isolated mouse microglial cells, bone-marrow-derived macrophages (BMDM) and peritoneal macrophages (PM) in a dose-dependent manner, with a maximum response at 6nM (Figure 6A-C). Adding Slit2 to both top and bottom chambers inhibited macrophage migration, indicating a chemotactic response (Supplemental Figure 9A-B).

**Figure 6.**
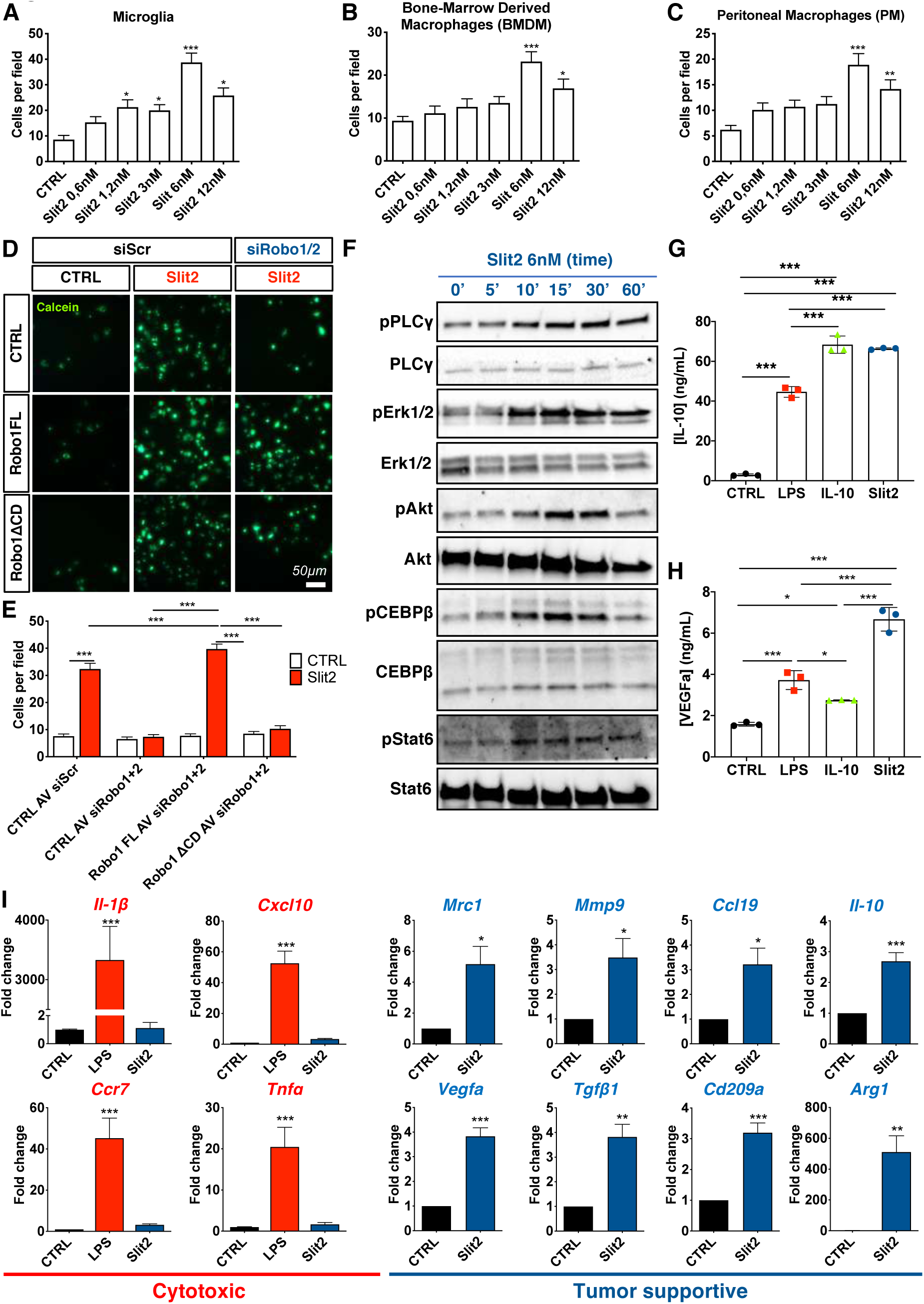
Slit 2 drives microglia and macrophage migration and tumor supportive polarization. **A-C.** Transwell assay of microglial cells (**A**), bone marrow derived macrophages (BMDM) (**B**) and peritoneal macrophages (PM) (**C**) in response to Slit2 or carrier (CTRL) in the bottom chamber (*n =* 4, One-way ANOVA). **D-E.** Transwell assay of RAW macrophages treated or not with Robo1/2 siRNA and infected with adenovirus encoding CTRL (GFP construct), Robo1FL or Robo1ΔCD constructs and stained with Calcein. **E.** Quantification of **(D)** (*n =* 3, Two-way ANOVA). **F**. Western blot analysis of Slit2 downstream signaling in cultured BMDM (*n =* 6). **G-H.** ELISA from conditioned medium from LPS, Il-10 or Slit2-treated BMDMs to quantify IL-10 (**G**) and VEGFa (**H**) (*n =* 3 independent cultures, Mann-Whitney U test)**. I.** qPCR analysis of BMDM cultures following Slit2 or LPS treatment (*n =* 4, One-Way ANOVA or Mann-Whitney U test). Data are presented as mean ± s.e.m. * *P* < 0.05, ** *P* < 0.01, *** *P* < 0.001

To determine if SLIT2 signaled through ROBO receptors to promote macrophage migration, we silenced *Robo1&2* in cultured RAW264.7 macrophages using siRNAs, which inhibited Slit2-induced macrophage migration (Figure 6D-E, Supplemental Figure 9C-D). Migration could be rescued by adenoviral-induced expression of a siRNA-resistant full-length rat Robo1 construct (Robo1FL) but not by a construct lacking the cytoplasmic signaling domain (Robo1ΔCD, Figure 6D-E).

To identify SLIT2 downstream signaling pathways in macrophages, we treated BMDM and microglial cells with 6nM Slit2, which led to PLCγ, Erk1/2 and Akt phosphorylation (Figure 6F, Supplemental Figure 9E-G, J). SLIT2 also induced phosphorylation of Stat6 and CEBPβ1 that polarize tumor-infiltrating macrophages towards a tumor supportive phenotype (63) (Figure 6F, Supplemental Figure 9H-J), suggesting that SLIT2 induced tumor-supportive gene expression changes.

Conditioned medium of Slit2-treated microglia and macrophages increased levels of IL-10 and VEGFa compared to cells not treated with Slit2 (Figure 6G-H, Supplemental Figure 9K-L). The expression of genes characteristic of a tumor supportive macrophage phenotype, including *Mrc1, Vegfa, Mmp9, Tgfβ1, Ccl19, Cd209a, Il-10* and *Arg1*, were all increased by Slit2 treatment, while cytotoxic response-related genes *Il-1β, Cxcl10, Ccr7* and *Tnfα* were unaffected by Slit2, but increased by LPS (Figure 6I). Slit2-induced gene expression changes were ROBO1&2-dependent, as shown by siRNA silencing of *Robo1/2,* which abrogated Slit2 induced changes in protein phosphorylation and gene expression (Supplemental Figure 10A-G).

### Slit2-Robo induced tumor-supportive macrophage/microglia polarization via PI3Kγ

Previous studies have shown that Stat6 and CEBPβ1 activation in TAMs occur downstream of PI3Kγ (63), leading us to ask if Slit2-Robo1&2 signaled upstream of PI3Kγ to induce macrophage tumor-supportive polarization. First, we observed Robo1 and PI3Kγ co-immunoprecipitation in BMDMs, which was enhanced after Slit2 treatment for 15 minutes (Figure 7A). Second, Slit2-induced BMDM migration was abrogated by pre-treatment with a specific PI3Kγ inhibitor IPI-549 (1μM) (Figure 7B). Third, Slit2-induced phospho-Stat6 nuclear translocation in cultured BMDMs was prevented by pre-treatment with IPI-549 (Figure 7C-D). Slit2-induced Akt and Stat6 phosphorylation (Supplemental Figure 11A), as well as IL-10 and VEGFa secretion in ELISA from BMDM conditioned medium were also reduced by PI3Kγ inhibition (Figure 7E-F). Finally, the Slit2-induced expression of genes characteristic of a tumor supportive macrophage phenotype (*Mrc1, Vegfa, Mmp9, Tgfβ1, Ccl19, Cd209a, Il-10* and *Arg1*) was disrupted by IPI-549 pretreatment, while LPS-induced cytotoxic response-related genes were unaffected in both BMDMs and microglial cells by PI3Kγ inhibition (Figure 7G, Supplemental Figure 11B).

**Figure 7.**
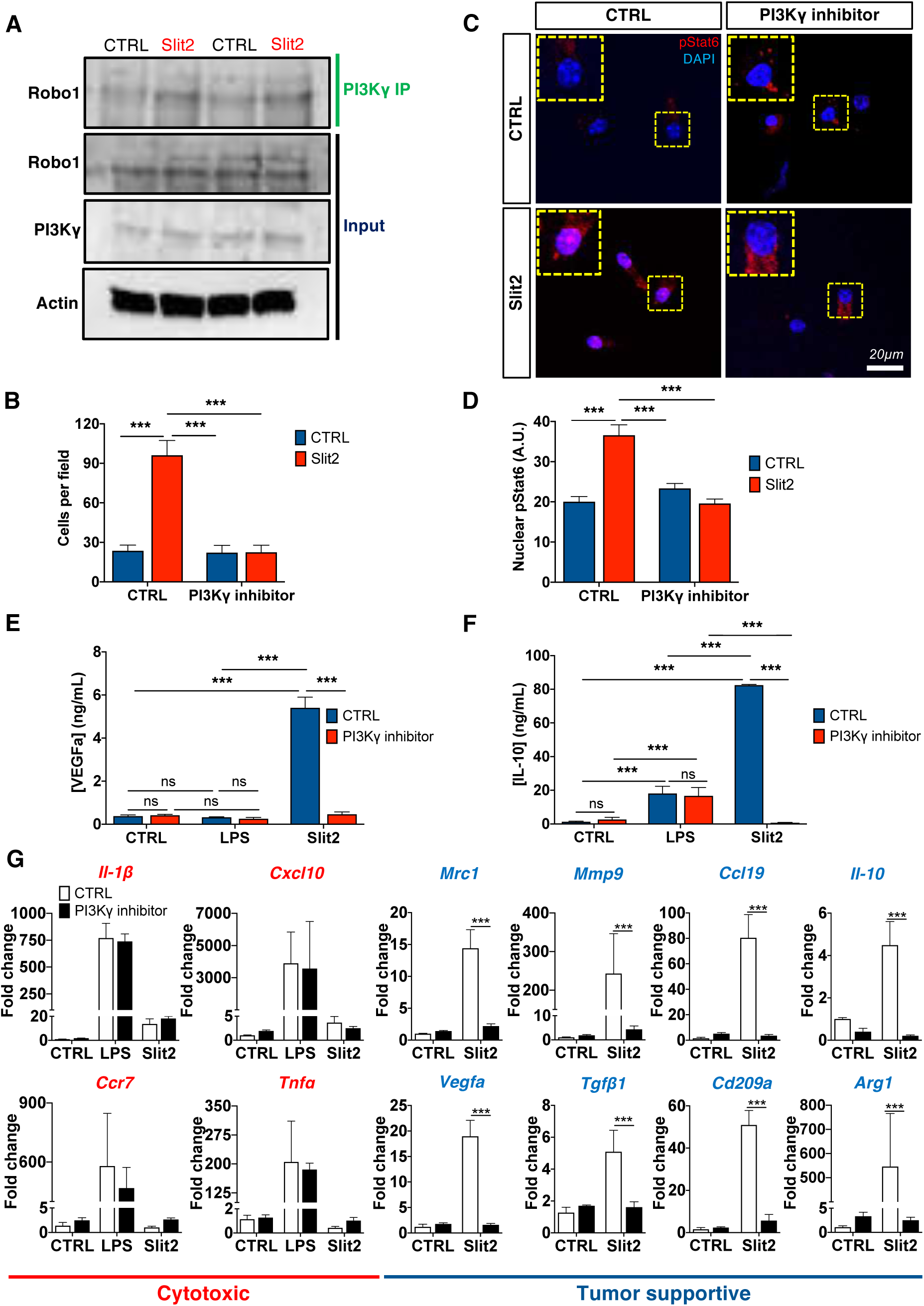
Slit2 driven microglia/macrophage polarization via PI3Kγ. **A.** PI3Kγ immunoprecipitation in BMDMs treated or not with Slit2 for 15 minutes and WB for Robo1 (*n =* 3 independent experiments). **B.** Transwell assay of BMDMs in response to Slit2 or carrier (CTRL) in the bottom chamber after pretreatment with vehicle control (DMSO) or PI3Kγ inhibitor IPI-549 (1uM). **C-D.** Phospho-Stat6 immunofluorescent staining of BMDMs treated or not with Slit2 and PI3Kγ inhibitor and quantification of nuclear pStat6 intensity (*n =* 4 independent cultures, 2-way ANOVA). **E-F.** ELISA from conditioned medium from LPS or Slit2-treated BMDMs with vehicle control (DMSO) or PI3Kγ inhibitor, to quantify IL-10 (**E**) and VEGFa (**F**) (*n =* 3 independent cultures, 2-way ANOVA)**. G**. qPCR analysis of BMDM cultures following Slit2 or LPS treatment with vehicle control or PI3Kγ inhibitor (*n =* 4 independent cultures, 2-way ANOVA). Data are presented as mean ± s.e.m. * *P* < 0.05, ** *P* < 0.01, *** *P* < 0.001

### Robo deficiency in TAMs inhibited glioma growth and vascular dysmorphia

To determine if SLIT2 signaling effects in macrophages were sufficient to drive the stromal response, we developed mice with genetic Robo receptor deletions in macrophages. To do so, we intercrossed Robo1^-/-^Robo2^fl/fl^ mice (28) with CSF-1R^CreERT2^ mice (64) on a ROSA^mT/mG^ background, generating Robo1^-/-^Robo2^fl/fl^CSF- 1R^CreERT2^ROSA^mT/mG^ mice (hereafter named iRoboMacKO mice). Littermate Robo1^+/-^ Robo2^+/fl^ CSF-1R^CreERT2^ROSA^mT/mG^ or Robo1^-/-^Robo2^fl/fl^ROSA^mT/mG^ mice were used as controls. Mice were implanted with CT-2A-BFP glioma cells and followed longitudinally during tumor growth. Tamoxifen injections to induce gene deletion were done every 3 days starting 7 days after tumor implantation, and induced robust gene deletion, assessed by qPCR of GFP^+^ macrophages extracted from the bone marrow (Supplemental Figure 12A-B).

MRI imaging and histological analysis 21 days after tumor implantation converged to show reduced tumor size in iRoboMacKO tumors when compared to controls (Figure 8 A-C). T1-weighted imaging after Gadolinium injection showed more homogeneous contrast signal in iRoboMacKO tumors, while control GBMs displayed predominantly peripheral and heterogenous contrast distribution, suggesting improved perfusion in iRoboMacKO tumors (Figure 8A). *In vivo* two-photon imaging revealed that blood vessels in iRoboMacKO tumors dilated less and remained more ramified when compared to controls (Figure 8D-F). Glut1^+^ hypoxic zones within the tumor mass were reduced in iRoboMacKO tumors, confirming improved perfusion when compared to controls (Figure 8G-H). Most of the Glut1 staining in iRoboMacKO tumors colocalized with Tomato^+^ blood vessels, attesting to the qualitative improvement of iRoboMacKO tumor vessels (Figure 8G).

**Figure 8.**
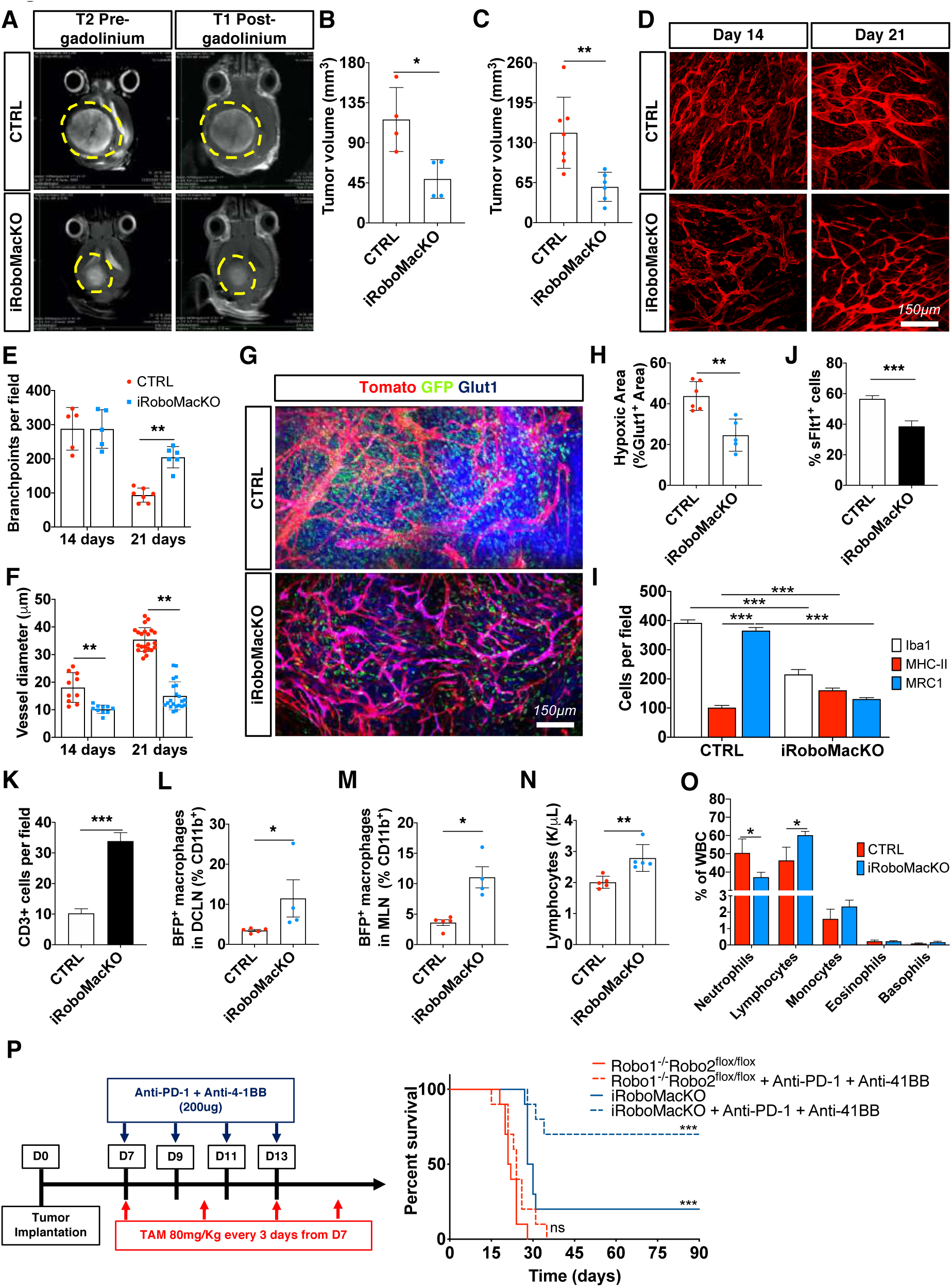
Macrophage-specific Robo1/2 KO normalizes the TME. **A.** T2-weighted pre-gadolinium and T1-weighted post-gadolinium MRI images of CTRL and iRoboMacKO mice 21 days after tumor spheroid implantation. **B-C.** Quantification of tumor size 21 days after tumor spheroid implantation on MRI images (**B**, *n =* 4 tumors per group, Mann-Whitney U test) and serial vibratome sections (**C**, *n =* 7 CTRL and 6 iRoboMacKO tumors, Mann-Whitney U test). **D-F.** *In vivo* two-photon images of tumor bearing mice in (**D**) and quantification of vessel diameter (**E**) and branchpoints (**F**) (*n =* 6 mice per group, One-way ANOVA). **G-H.** Glut1 (blue) immunohistochemistry on day 21 tumor bearing CTRL and iRoboMacKO mice (**G**), and quantification of hypoxic areas in the tumor (**H**) (*n =* 6 CTRL and 5 iRoboMacKO tumors, Mann-Whitney U test). **I-K.** Quantification of immunohistochemistry on sections of day 21 CT-2A CTRL and iRoboMacKO tumors for F4/80, MHC-II and MRC1^+^ cells (**I**), VEGFa-expressing (sFLT1^+^) GFP^+^ cells (**J**), and total TALs (CD3^+^) (**K**) (*n =* 6 CTRL and 5 iRoboMacKO tumors, 2-way ANOVA or Mann-Whitney U test). **L-M.** FACS analysis of deep cervical and mandibular lymph nodes (DCLN and MLN, respectively) from day 21 CTRL and iRoboMacKO tumor-bearing mice (*n =* 5 CTRL and 4 iRoboMacKO mice; Mann-Whitney U test). **N-O.** Lymphocyte counts (**N**) and differential WBC counts (**O**) from peripheral blood of day 21 CTRL and iRoboMacKO tumor-bearing mice (*n =* 5 mice/group; Mann-Whitney U test). **P.** 8-week-old mice were engrafted with CT-2A BFP and gene deletion was achieved by 80mg/kg Tamoxifen intraperitoneal injection every 3 days starting 7 days after tumor implantation. Robo1^-/-^Robo2^flox/flox^ and iRoboMacKO mice were randomly assigned to vehicle or Anti-PD-1 + Anti-4-1BB treatment (200ug per dose on days 7, 9, 11 and 13 after tumor implantation) (*n =* 10/11 mice per group, O.S.= 21.5 days for Robo1^-/-^Robo2^flox/flox^, 24 days for Robo1^-/-^Robo2^flox/flox^ + Anti-PD-1 + Anti-4-1BB, 29 days for iRoboMacKO and Undetermined for iRoboMacKO + Anti-PD-1 + Anti-4-1BB; Multiple comparisons log-rank test). Data are presented as mean ± s.e.m. * *P* < 0.05, ** *P* < 0.01, *** *P* < 0.001

Compared to controls, iRoboMacKO displayed reduced overall numbers of intra-tumor Iba1^+^ myeloid cells, with a significant increase of cytotoxic MHCII^+^ cells and a reduction in tumor-supportive MRC1^+^ cells (Figure 8I, Supplemental Figure 12D). Soluble Flt1 binding was reduced in iRoboMacKO tumors (Figure 8J, Supplemental Figure 12D), and Robo1/2-deleted macrophages extracted from the bone marrow of tumor-bearing mice showed decreased *Vegfa* expression (Supplemental Figure 12C).

T cell infiltration was increased in iRoboMacKO tumors (Figure 8K, Supplemental Figure 12D), suggesting that SLIT-ROBO signaling inhibition in macrophages was sufficient to shift the GBM microenvironment towards a cytotoxic, T cell enriched phenotype. This effect could be due to increased circulation of antigen-presenting cells (APCs) to the tumor draining lymph-nodes, where they can activate anti-tumor T cell responses (59). Analysis of glioma-draining deep cervical and mandibular lymph nodes (DCLN and MLN, respectively) for the presence of BFP tumor antigen in immune cells revealed an important increase in CD11b^+^BFP^+^ cells in both deep cervical (DCLN) and mandibular lymph nodes (MLN) of iRoboMacKO tumors when compared to controls (Figure 8L-M, Supplemental Figure 12E).

Lymphocyte sequestration in the bone marrow contributes to the T cell-depleted TME and failure of currently available immunotherapy (11). iRoboMacKO mice had significantly increased lymphocyte counts in peripheral blood 21 days after tumor implantation (Figure 8N). Given that total white blood cell (WBC) count was not changed (Supplemental Figure 12F), tumor-bearing iRoboMacKO mice shifted to a predominance of lymphocytes over neutrophils in the blood stream (Figure 8O), revealing a reduction in the systemic immunosuppression after macrophage-specific Robo1&2KO.

Given the profound changes observed in the TME observed, we next tested if macrophage-specific Robo1&2 deletion was sufficient to prolong survival and sensitivity to checkpoint inhibitor therapy. Indeed, macrophage-specific Robo1/2 knockout increased tumor-bearing mice survival (Figure 8P, O.S., 21.5 days for Robo1^-/-^Robo2^fl/fl^, 29 days for iRoboMacKO), and survival benefit was further increased by immune checkpoint inhibitors, with 70% of the iRoboMacKO mice alive after 100 days (Figure 8P, O.S., 24 days for Robo1^-/-^Robo2^fl/fl^ + Anti-PD-1+Anti-4-1BB, Undefined for iRoboMacKO + Anti-PD-1+Anti-4-1BB).

In contrast to macrophage Robo depletion, T cell depletion using anti-CD3 145-2C11 antibodies (65) did not induce significant changes of blood vessels or TAMs in the GBM microenvironment (Supplemental Figure 13).

### Systemic SLIT2 inhibition alleviated GBM immunosuppression

We reasoned that systemic administration of a SLIT2 ligand trap protein (Robo1Fc) might be efficient in a therapeutic setting. Mice with established shCTRL CT2A tumors were intravenously injected 5 times with 2.5 mg/kg of Robo1Fc every second day starting from day 7 after tumor implantation and analyzed at day 23 (Figure 9A). Control mice received injections of human control IgG1 Fc fragment. Robo1Fc treatment reduced Slit2 serum levels, as attested by Slit2 ELISA on days 14 and 21 after tumor implantation (Figure 9B). Mice treated with Robo1Fc exhibited a pronounced tumor growth reduction compared to control Fc-treated tumors (Figure 9C-D). MRI analysis 21 days after tumor implantation showed that tumor size was reduced and that tumor perfusion was improved, as seen by the more homogeneous gadolinium uptake in Robo1Fc-treated tumors compared to controls (Supplemental Figure 14A-B). *In vivo* imaging demonstrated that Robo1Fc treatment reduced vascular dysmorphia (Figure 9E-G) and reduced Glut1^+^ hypoxic zones within the tumor mass (Figure 9H, Supplemental Figure 14C). Robo1Fc treatment changed immune cell infiltration and reduced overall numbers of intra-tumoral F4/80^+^ cells, with a significant increase of cytotoxic MHCII^+^ cells and a reduction of tumor-supportive MRC1^+^ cells compared to controls (Figure 9I, Supplemental Figure 14D). Soluble Flt1 binding was reduced in Robo1Fc treated tumors (Figure 9J, Supplemental Figure 14D), while T cell infiltration was increased compared to controls (Figure 9K, Supplemental Figure 14D).

**Figure 9.**
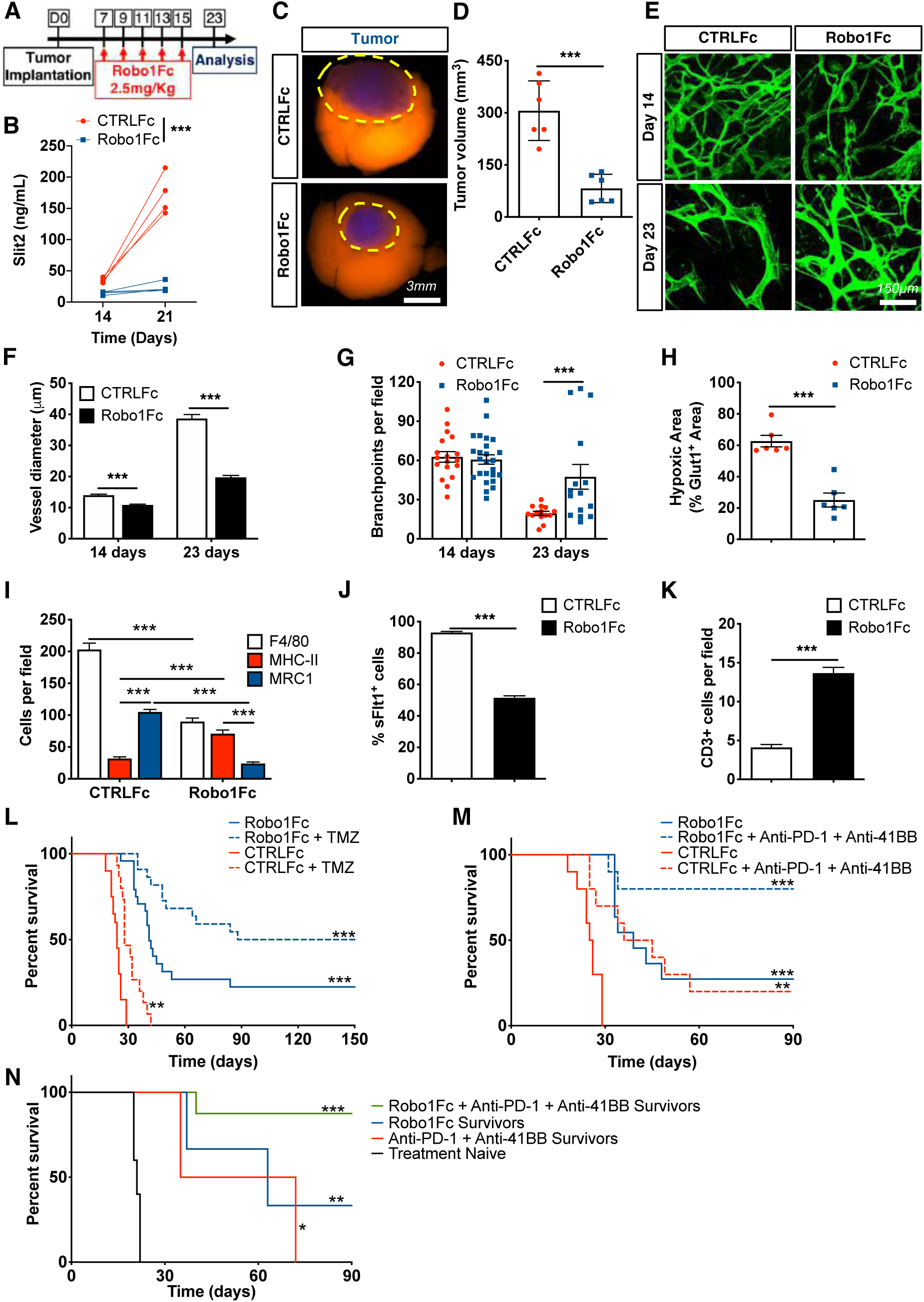
Robo1Fc treatment limits glioma growth by shifting the microenvironment. **A.** Experimental design: 8-week-old mice were engrafted with CT-2A spheroids and randomly assigned to CTRLFc or Robo1Fc treatment (2.5 mg/kg) every other day between day 7 and day 15 after tumor implantation. **B.** ELISA dosage of serum Slit2 in CTRLFc- and Robo1Fc-treated mice at days 14 and 21 (*n =* 4 mice per group, Two-Way ANOVA). **C.** Representative CTRLFc- and Robo1Fc-treated tumors at 23 days. **D**. quantification of **(C)** (*n =* 6, Student’s T test). **E**. *In vivo* two-photon images of day 23 CTRLFc and Robo1Fc treated CT-2A tumors. **F-G.** Quantification of vessel diameter (**F**) and branchpoints (**G**) from (**E**) (*n =* 6 mice per group, One-way ANOVA). **H**. Quantification of hypoxic areas (Glut1^+^) on stained tumor sections of day 23 tumor-bearing mice treated with CTRLFc or Robo1Fc (*n =* 6 mice per group, Mann-Whitney U test). **I-K**. Quantification of F4/80, MHC-II and MRC1 (**I**), soluble-Flt1 binding (**J**) and CD3 (**K**) immunostaining of day 23 tumor-bearing mice treated with CTRLFc or Robo1Fc (*n =* 6 mice per group, Two-way ANOVA or Student’s t-test). **L.** 8-week-old wild-type mice were injected with CT-2A cells and assigned randomly to one of the following treatments: CTRLFc + vehicle (*n =* 20), CTRLFc + TMZ (40mg/kg on days 7, 11, 15 and 19 after tumor implantation) (*n =* 15), Robo1Fc + vehicle (*n =* 24) or Robo1Fc + TMZ (*n =* 22, Multiple comparisons Mantel–Cox log-rank; O.S., CTRLFc: 24 days; CTRLFc + TMZ: 28 days; Robo1Fc: 41 days; Robo1Fc + TMZ: 119 days). **M.** 8-week-old wild-type mice were injected with CT-2A cells and assigned randomly to one of the following treatments: CTRLFc + vehicle, CTRLFc + Anti-PD-1 + Anti-4-1BB treatment (200ug of each on days 7, 9, 11 and 13 after tumor implantation), Robo1Fc + vehicle or Robo1Fc + Anti-PD-1 + Anti-4-1BB treatment (*n =* 10/11 mice per group; O.S., CTRLFc: 25.5 days; CTRLFc + Anti-PD-1 + Anti-4-1BB treatment: 40 days; Robo1Fc: 39 days; Robo1Fc + Anti-PD-1 + Anti-4-1BB treatment: Undetermined; Multiple comparisons log-rank test). **N.** 90 days after tumor implantation, surviving mice from Figure 9M (*n =* 2 Anti-PD-1 + Anti-4-1BB survivors, 3 Robo1Fc survivors and 8 Robo1Fc + Anti-PD-1 + Anti-4-1BB survivors) or 8-weeks-old tumor naïve mice (*n =* 10 mice) were re-challenged by implantation of CT-2A cells in the contralateral hemisphere from the first injection (O.S., Naïve mice: 21 days; CTRLFc + Anti-PD-1 + Anti-4-1BB survivors: 53.5 days; Robo1Fc survivors: 63 days; Robo1Fc + Anti-PD-1 + Anti-4-1BB treatment: Undetermined; Multiple comparisons log-rank test). Data are presented as mean ± s.e.m. * *P* < 0.05, ** *P* < 0.01, *** *P* < 0.001.

Analysis of glioma-draining DCLN and MLN showed an increased presence of GFP tumor antigen in APCs (CD45^+^CD11b^+^Ly6G^-^) of Robo1Fc-treated mice when compared to CTRLFc treated ones (Supplemental Figure 14E-G), as we observed in iRoboMacKO mice. Finally, Robo1Fc-treated mice also had significantly increased total WBC and lymphocyte counts in peripheral blood 21 days after tumor implantation, and we observed a shift to a predominance of Lymphocytes over Neutrophils in the blood stream of Robo1Fc treated mice, while other white blood cell counts were unchanged (Supplemental Figure 14H-J).

Five injections of Robo1Fc protein during early stages of tumor progression were sufficient to significantly extend survival of tumor-bearing mice and 25% of the treated mice survived 150 days after implantation (Figure 9L, O.S., 24 days for CTRLFc and 41 days for Robo1Fc). Combining Robo1Fc with TMZ further increased this survival benefit, with 45% of the mice surviving 150 days after implantation (Figure 9L, O.S., 28 days for CTRLFc + TMZ and 119 days for Robo1Fc + TMZ). Combining Robo1Fc with Anti-PD-1 and Anti-4-1BB antibodies further improved anti-tumor responses, with 80% of the mice surviving 90 days after tumor implantation (Figure 9M, O.S., 40.5 days for CTRLFc + Anti-PD-1 + Anti-4-1BB and Undefined for Robo1Fc + Anti-PD-1 + Anti-4-1BB). Mice that survived the immunotherapy were rechallenged by a novel tumor injection in the contralateral hemisphere. Mice that survived after treatment with Robo1Fc and checkpoint inhibitors had the best long-term survival after tumor rechallenge, with more than 80% of mice alive 90 days after tumor re-injection (Figure 9N, O.S., 22 days for naïve mice, 53.5 days for Anti-PD-1 + Anti-4-1BB survivors, 63 days for Robo1Fc survivors and Undefined days for Robo1Fc + Anti-PD-1 + Anti-4-1BB survivors)

## Discussion

Collectively, our data showed that GBM-derived SLIT2 signaled through ROBO1/2 in TAMs, which resulted in an impairment of anti-tumor immunity and the induction of vascular dysmorphia in the TME. SLIT2-ROBO1&2 signaling is therefore a novel immune evasion mechanism in the TME, and inhibiting this pathway in TAMs could sensitize GBM to immune checkpoint inhibitors, and add to the therapeutic arsenal against GBM.

The main findings of our study can be summarized as follows: we showed that *SLIT2* expression levels correlated with tumor aggressiveness, poor prognosis and immunosuppression in a variety of glioma patient cohorts. In particular, low-grade glioma patients with low *SLIT2* expression levels had a significantly prolonged survival when compared to those with higher *SLIT2* expression, suggesting that SLIT2 may be a useful prognostic marker for glioma patients.

Our data suggest that GBM tumor cells are a major source of SLIT2. *SLIT2* expression in human GBM tumors was highest in the tumor cell compartment, and *Slit2* knockdown in two murine GBM cells lines and in a human PDX model decreased tumor growth, while SLIT2 overexpression in CT2A cells enhanced murine GBM tumor growth. SLIT2 from other cell compartments could also affect GBM growth, but since genetic *SLIT2* inhibition in tumor cells and systemic SLIT2 inhibition had similar effects in our mouse models, it is likely that tumor cell SLIT2 plays an important role in GBM. In further support of this idea, silencing tumor cell derived *SLIT2* in human PDX GBM tumors reduced tumor development *in vivo*.

We observed that SLIT2 acted on different cell types within GBM. First, both human and mouse GBM tumor cells expressed ROBO1&2 receptors. *SLIT2* knockdown did not affect tumor cell proliferation or survival, and SLIT2 did not attract tumor cells in transwell assays *in vitro.* However, *SLIT2* knockdown decreased tumor cell migration towards a serum gradient in transwell chambers and reduced spheroid invasion in fibrin gels, and patient-derived shSLIT2 GBM cells implanted in Nude mice decreased invasiveness compared to shCTRL. These results are consistent with SLIT2-ROBO signaling driving pro-invasive GBM tumor cell behavior in both mice and patient-derived models. Our data contrast early studies with commercial human GBM cell lines where SLIT2-ROBO1 signaling inhibited migration (51, 52), but they support and extend s studies using murine GBM models (55) and patient-derived tumor spheres and xenograft models (54), which showed that SLIT2-ROBO1 signaling in tumor cells promotes tumor invasiveness.

In addition to the tumor cells themselves, SLIT2 exerted major effects in the TME, and remarkably these changes appeared centered around ROBO1&2 signaling in macrophages/microglial cells. We found that in the tumor context, ROBO1&2 signaling inhibition in macrophages was sufficient to recapitulate all major aspects of tumor cell SLIT2 manipulation, or systemic SLIT2 inhibition with a ligand trap, and shifted the entire TME towards a normalized and cytotoxic phenotype. We identified three TME cell types that responded to tumor cell SLIT2, namely TAMs, endothelial cells of blood vessels and T cells. Genetic inhibition of *Robo* signaling in macrophages reduced macrophage recruitment to the TME, prevented phenotypic conversion into tumor-supportive macrophages, normalized tumor vasculature and induced T-cell based anti-tumor responses. It remains possible that cell-autonomous *Robo* signaling in endothelial cells, which induces angiogenesis (28, 31), or T-cell *Robo* signaling contribute to the observed effects in GBM, but clearly macrophage *Robo* signaling appeared dominant.

Mechanistically, SLIT2-mediated TAM migration and polarization were ROBO1&2 dependent and mediated by PI3Kγ signaling. PI3Kγ signaling inhibition has been previously shown to prevent TAM polarization and tumor progression in different cancer models (63), and this mechanism could be conserved in GBM TAMs. PI3Kγ is traditionally activated by G-protein coupled receptors (GPCRs) or Receptor tyrosine kinases (RTKs), therefore it remains to be established how ROBO activates PI3Kγ mechanistically, via NCK-SOS activation of RAS or other small GTPases that can activate PI3Kγ (66–68). Another possibility is that PI3Kγ activation downstream of ROBO receptors depends on the co-activation of other RTKs or GPCRs and their endocytosis.

TAMs are the most abundant cells in the GBM microenvironment, and are known to contribute to immunosuppression in the TME (7–9, 69) and dysmorphic angiogenesis (16, 70–72). Hence, TAMs are key players in the development of resistance to anticancer therapies (17, 73–76). Several attempts have been made to target TAM signaling for GBM treatment, including manipulation of VEGFa and angiopoietins, Neuropilin1 (77, 78), CD-47 or CSF-1R (79–81). Combined VEGF/Angiopoietin inhibition led to vascular normalization and cytotoxic TAM polarization, but did not change T cell infiltration or activation profile (82, 83). CD47 inhibition prolonged GBM-bearing mice survival due to increased phagocytosis capacity and cytotoxic TAM polarization, but did not affect other components of the GBM microenvironment (84). CSF-1R inhibition did not change TAM production of pro-angiogenic molecules such as VEGFa and therefore did not lead to vascular normalization in GBM (16, 79, 81). Hence, these strategies changed the TAM component of the GBM microenvironment, but they did not induce the profound changes in angiogenesis and T cell response achieved by SLIT2 inhibition. Systemic SLIT2 inhibition via intravenous injection of a SLIT2 ligand trap could be optimized and translated into clinical practice to combat GBM in human patients, especially those with high levels of *SLIT2* expression.

## Methods

### Bioinformatic analysis

For ‘The Cancer Genome Atlas’ (TCGA) dataset, RNAseqV2 normalized data (level 3, log2(x+1) transformed RSEM normalized count, version 2017-10-13) of 151 primary glioblastoma multiforme patients (TCGA Glioblastoma (GBM)) and associated molecular GBM subtypes and clinical data were downloaded from the cBioPortal website datapages (https://www.cbioportal.org/study/summary?id=gbm_tcga). The cohort was split into 3 groups of patients defined by the level of their expression. Overall survival (in months) was used to estimate survival distributions using the Kaplan–Meier method and the distributions were compared using the log-rank test.

### Patient Samples

Frozen tumors samples were obtained from 25 patients after informed consent and approval by UZ Leuven ethical committee for the Brain-Tumor-Imm-2014 study; and tumor RNA was obtained from 104 patients of the Pitié-Salpêtrière tumor bank Onconeurotek.

RNA was purified from liquid nitrogen frozen tissue samples using RNeasy-kit (Qiagen). 0.5μg of RNA were reverse transcribed using SuperScript IV Reverse Transcriptase and Random Primers (Invitrogen) for qPCR reactions.

### QPCR reactions

Real-time quantitative PCR (qPCR) reactions were performed in duplicate using the MyIQ real-time PCR system (Bio-Rad), with iQ SYBR Green Supermix (Bio-Rad) and QuantiTect qPCR primers (Qiagen). Each reaction contained 10 ng of cDNA and 250 nM forward and reverse primers. Fold changes were calculated using the comparative CT method.

### Single-cell RNA sequencing analysis

We downloaded the following published datasets for single cell RNA-seq analysis from GEO: GSE138794, GSE131928, and GSE84465 (85–87). Gene expression matrices were combined and were visualized using the Seurat v3 (88) package in R. Based on the ElbowPlot function, we chose around 43 principal components for UMAP driven visualizations. Markers for each cluster were defined from a combination of literature knowledge and the FindMarkers function in Seurat. For removal of batch effects between different datasets, we used the harmony package (89).

### Cell lines

RAW264.7 mouse macrophages, CT-2A and GL261 glioma cells were cultured in DMEM Gluta-MAX (Gibco) supplemented with 10% FBS (Gibco), 1% penicillin/streptomycin (Gibco) until a maximum of 10 passages. Glioma spheroids were obtained by seeding the glioma cells for 48 h on non-adherent culture dishes.

### Animal procedures and glioma implantation

All *in vivo* experiments were conducted in accordance to the European Community for experimental animal use guidelines (L358-86/609EEC) with protocols approved by the Ethical Committee of INSERM (n°MESRI23570 and #17503 2018111214011311 v5). Animals were housed with free access to food and water in a 12h light/dark cycle. For survival experiments, mice were euthanized if they exhibited signs of neurological morbidity or if they lost > 20% of their body weight. C57bl6J and ROSA^mT/mG^ mice were used for survival and live imaging experiments, respectively. For generation of macrophage-specific Robo1/2 KO, Robo1^-/-^Robo2^fl/fl^ mice (28) were bred with CSF1-R-CreERT2, ROSA^mT/mG^ mice (64). Gene deletion was induced by injections of 80mg/Kg of tamoxifen every 2 days starting 7 days after tumor implantation and was verified on GFP^+^ bone marrow monocytes/macrophages.

### Murine Glioma model

Craniotomy and glioblastoma spheroid implantation were done as previously described (16). Briefly, a 5-mm circle was drilled between sutures of the skull on ketamine/xylazine anesthetized mice. A 250-μm diameter CT-2A or GL261 spheroid was injected in the cortex and sealed with a glass coverslip. For survival experiments involving PD-1 and 4-1BB inhibition, tumor cells were inoculated as cell suspension in the mice striatum instead as cortical spheroids as previously described (59). Following intramuscular administration of analgesic (buprenorphine 1 mg/kg), mice were placed in a heated cage until full recovery.

For Temozolomide (Sigma) treatment, mice were injected intraperitoneally with 40mg/kg in 0,2 mL of PBS at days 7, 11, 15 and 19 after tumor implantation. For anti-PD1 (clone RMP1-14, BioXCell) and anti-4-1BB (clone LOB12.3, BioXCell) treatment, glioma-bearing mice were injected intraperitoneally with 0.2 mg of antibodies on days 7, 9, 11 and 13 after tumor implantation.

For anti-CD3 145-2C11 monoclonal antibody (BioXCell) treatment, 7-day glioma-bearing mice were injected intravenously with 0.2 mg of antibodies every 3 days and analyzed at 23 days of tumor growth.

For Robo1Fc (R&D Systems) treatment, 1-week growth glioma-bearing mice were injected intravenously with 2.5 mg/kg of Robo1Fc or human control IgG1 Fc fragment at days 7, 9, 11, 13 and 15 after tumor implantation. For this experiment, 6 different series of mice were implanted and treated: 2 for tumor volume measurement and histological analysis and 4 for survival analysis.

At the defined time points, blood samples were obtained by retro-orbital bleeding with EDTA-coated capillaries and complete blood cell counts were obtained with a HemaVet (Drew Scientific). 21 or 23 days after tumor implantation, anesthetized mice were transcardially perfused with 2% PFA solution. The mouse brain was harvested and fixed overnight in 4% PFA at 4°C. For immunohistochemistry, brains were washed with PBS and sectioned with a vibratome (200um-400μm sections). Tumor volume was measured on serial 400μm sections of the whole tumor under a stereo-microscope using Leica software according to Cavalieri’s principle.

### Slit2 shRNA knockdown and overexpression

CT-2A and GL261 glioma cell lines were infected with Slit2 mouse shRNA lentiviral particles (Locus ID 20563, Origene TL511128V) in accordance with the manufacturer’s instructions. After infection, cells were polyclonally selected by Puromycin and GFP^+^ cells were sorted by FACS. Slit2 knockdown was verified by qPCR and Western Blot analysis, and cells were implanted after a maximum of 5 passages. For Slit2 re-expression, shSlit2 CT-2A cells were infected with SLIT2 (NM_004787) Human Tagged ORF Clone Lentiviral Particle (Origene) in accordance with manufacturer’s instructions. Cells were implanted after a maximum of 3 passages.

### FDG PET-CT Imaging

Mice were fasted overnight with free access to water. Mice were anesthetized with isoflurane, weighed and glycemia was measured in blood drawn from the caudal ventral artery using an Accu-Chek® Aviva Nano A (Accu-Chek, France). A 26G needle catheter (Fischer Scientific, France) connected to a 5cm polyethylene tubing (Tygon Microbore Tubing, 0.010” x 0.030”OD; Fisher Scientific, France) was inserted in the caudal vein for radiotracer injection. 9.2±1.5 MBq of 2’-deoxy-2’-[18F]fluoro-D-glucose (FDG; Advanced Applied Applications, France) in 0.2mL saline was injected via the catheter. Mice were left on a warming pad for 30 min and then installed into the PET-CT dedicated bed. Respiration and body temperature were registered. Body temperature was maintained at 34±2 °C and anesthesia was controlled on the breathing rate throughout the entire PET-CT examination. CT was acquired in a PET-CT scanner (nanoScan PET-CT; Mediso Medical Imaging Systems, Hungary) using the following acquisition parameters: semi-circular mode, 50kV tension, 720 projections full scan, 300ms per projection, binning 1:4. CT projections were reconstructed by filtered retro-projection (filter: Cosine; Cutoff: 100%) using the software Nucline 3.00.010.0000 (Mediso Medical Imaging Systems, Hungary). 55 min post tracer injection, PET data were collected for 10 min in list mode and binned using a 5ns time window, with a 400-600keV energy window and a 1:5 coincidence mode. Data were reconstructed using the Tera-Tomo reconstruction engine (3D-OSEM based manufactured customized algorithm) with expectation maximization iterations, scatter and attenuation correction. Volumes-of-interest (VOI) were delineated on the tumor and the contralateral brain on PET/CT fusion slices using the PMOD software package (PMOD Technologies Ltd, Zürich, Switzerland). Total FDG uptake was estimated as the product from the volume by the mean uptake of the segmented region.

### Live imaging

For multiphoton excitation of endogenous fluorophores in experimental gliomas, we used a Leica SP8 DIVE *in vivo* imaging system equipped with 4tune spectral external hybrid detectors and an InSightX3 laser (SpectraPhysics). The microscope was equipped with in house designed mouse holding platform for intravital imaging (stereotactic frame, Narishige; gas anesthesia and body temperature monitoring/control, Minerve). Acquisition of ROSA^mTmG^ reporter mice was performed at 1040-nm fixed wavelength. GFP signal from genetically modified tumor cells was acquired at 925-nm wavelength. Alexa Fluor 647 coupled Dextran was acquired at 1200-nm wavelength.

### Flow-cytometric staining of tumor-infiltrating immune cells

Day 21 CT-2A shCTRL and shSlit2 tumors were harvested, dissociated and incubated with anti-CD45 Alexa Fluor 594 (R&D Systems) or BUV805 (BD), anti-CD11b BV450 (BD), anti-Ly6G PerCP/Cy5.5 (BD), anti-Ly6C APC/Cy7 (BD), anti-F4/80 PE (BD), anti-CD11c (APC), anti-MHCII PE/Cy7 (Biolegend), anti-MRC1 BV711 (Biolegend), anti-CD3 PE/Cy5 (Biolegend), anti-CD19 PE/Texas Red (BD), anti-CD4 PE (Biolegend), and anti-CD8 PerCP/Cy5.5 (Biolegend). As a control, cells were stained with the appropriate isotype control. Data acquisition was performed on the BD LSRFortessa X20 and analysis was performed with FlowJo_V10.

### Slit2 ELISA

Slit2 concentrations in mice serum were determined by the sandwich ELISA method with the DuoSet ELISA Ancillary Reagent Kit 2 (R&D Systems) according to the manufacturer’s instructions, using serum samples obtained either from healthy mice or from tumor-bearing mice. Rat anti-Human/Mice Slit2 monoclonal antibodies (Clone 710305, R&D Systems) was used as capture antibody at a concentration of 1ug/mL and sheep anti-mouse Slit2 polyclonal antibody was used as detection antibody at a concentration of 400ng/mL HRP-linked anti-sheep secondary antibodies (1:1000) were used for revelation.

### MRI

Magnetic resonance imaging (MRI) was performed 21 days after tumor implantation in mice under Isofluorane anesthesia (2 to 2,5% mixed in ambient air) in a 4.7-T magnetic resonance scanner (Bruker BioSpec 47/40USR). Brain images were obtained using a Fast-Spin-Echo (FSE) T2 weighted (TE/TR: 15/2000 ms; matrix: 128×128; slice thickness: 1 mm; with no gap; 12 averages) and a Spin-Echo (SE) T1 weighted (TE/TR: 15/250 ms; matrix: 128×128; slice thickness: 1 mm; with no gap; 12 averages) sequences in axial and coronal planes. T1 weighted images were acquired before and T2 weighted images after intraperitoneal injection of gadoteric acid (200uL, 0.01mmol/mL, 0.05 M/Kg).

### siRNA transfection

Robo1, Robo2 and control siRNAs were purchased from Origene. We transfected RAW 264.7 macrophages with 10nM final siRNA concentration using siTran1.0 transfection reagent (Origene), according to the manufacturer’s instructions. Cells were used for experiments 72 h after transfection. For qPCR experiments, RNAs were purified using RNeasy-kit (Qiagen). 500 or 750 ng of RNA were reverse transcribed using SuperScript II Reverse Transcriptase and Random Primers (Invitrogen). Quantitative PCR were assayed as described for patient samples.

For adenoviral Robo1 rescue, we used previously described methods (28, 31).

### Transwell Migration Assay

For chemotactic migration assays with 8.0μm Polycarbonate Membrane Transwell inserts (Corning Inc), 20.000 primary cells were plated in 125 μL of serum-free DMEM medium on the top chambers. When stated, 1.000ng/mL of rmSlit2 was also added to the top chambers. Then, bottom chambers were filled with 500 μL of serum-free DMEM with chemoattractants (R&D Systems). Cells were cultured overnight at 37°C and 5% CO_2_, then incubated for 30 minutes with Calcein AM (Invitrogen) to stain live cells. Then the wells were washed and 10 pictures per well were acquired at 10x magnification using a Leica DMIRB inverted epifluorescence microscope. Migrated cells per field were counted using ImageJ software.

For Transwell migration assay in direction to tumor cells, 30.000 tumor cells were plated in the bottom chamber and starved in 500uL of serum-free media for 8 hours before plating cells on the top chamber.

### Western blot analysis

After siRNA transfection and/or treatments, cells were lysed in RIPA lysis buffer including phosphatase and protease inhibitors (Invitrogen). Equal amounts of proteins were separated on 4–15% Criterion precast gel (Bio-rad) and transferred on nitrocellulose membrane with Transblot Turbo (Bio-rad). Then membranes were blocked in 5% non-fat milk in TBS-T for 30 minutes at room temperature and incubated with primary antibodies against Robo1 (R&D Systems, 1:500), Robo2 (R&D Systems, 1:500), Actin (Sigma, 1:4000), anti-phospo p44/42 MAP kinase (phospho-ERK, Cell Signaling, 1:1000), anti-p44/42 MAP kinase (total ERK, Cell Signaling, 1:1000), anti-pAkt Ser473 (Cell Signaling, 1:1000), anti-Akt (Cell Signaling, 1:1000), anti-pPLCγ Ser 1248 (Cell Signaling, 1:1000), anti-PLCγ (4511T, Cell Signaling, 1:1000) overnight at 4°C under agitation. After washing with TBS-T membranes were incubated with proper HRP-conjugated secondary antibodies for 3 hours at room temperature under agitation. Western blots were developed with chemiluminescence HRP substrate (Bio-rad) on a Luminescent image analyser, ChemiDoc XRS+ (Bio-rad).

### Statistical analysis and quantification

For continuous variables, data are presented as mean ± s.e.m. Between-group comparisons used the Mann–Whitney U-test or t-test depending on the sample size for continuous variables. In cases where more than two groups were compared, one-way ANOVA test was performed, followed by Turkey’s multiple comparison test, and results were considered significantly different if *P* < 0.05. For comparisons involving grouped data, two-way ANOVA test was performed, followed by Turkey’s multiple comparison test, and results were considered significantly different if *P* < 0.05.

For survival experiment, log-rank (Mantel–Cox) tests and multiple comparison tests were performed. A two-tailed value of *P* < 0.05 was considered statistically significant. All the analyses were performed using Prism 6.0 software (GraphPad).

For mice *in vivo* imaging quantification, four to nine fields per animal were pictured in the tumor center and blood vessel caliber, branching and vessel perfusion were analyzed using Fiji software.

For mice *ex vivo* imaging quantification, five fields per individual were pictured in the tumor center and number of macrophages, overlapping stainings, hypoxic area and tumor double-strand DNA damages were quantified using Fiji software.

## ACKNOWLEDGEMENTS

This study was supported by the Fondation ARC pour la Recherche sur le Cancer–Institut National du Cancer (ARC-INCa, A.E., JLT and A.B.), ERC (grant agreement No. 834161 to AE) and NHLBI (1R01HLI125811 to AE.). L.H.G. was founded by fellowships of the ‘Coordenação de Aperfeiçoamento de Pessoal de Nível Superior’ (CAPES) and the European Society of Cardiology (ESC Basic Research Fellowship). We thank the Paris Cardiovascular Research Center (PARCC) Flow and Image Cytometry facility (Drs. Camille Brunaud and Camille Knosp), FDG PET and MRI imaging platform (France Life Imaging network (grant ANR-11-INBS-0006) and ‘Infrastructures Biologie Santé’ (IBISA), and multiphoton imaging platform (Leducq Foundation RETP: Visualisation des pathologies vasculaires). We thank Dr. Holger Gerhardt for providing GL261, CT-2A and RAW264.7 cells and CSF1-R-CreERT2 mice and Dr. Alain Chedotal for providing Robo1^-/-^Robo2^fl/fl^ mice used in this study.

## AUTHOR CONTRIBUTIONS

LHG, AE, JLT and TM conceived and designed the project. LHG, YX, LJ, LPF, RR, NM, MV, CK, CL, TD, JR, TV and TM performed experiments, collected and analyzed data. JD, LS, SDV and MS collected patient samples and provided clinical data for the study. BT, AB, AI, RL, FRSL contributed to study design and reviewed and revised the manuscript. FRSL, AE and TM supervised the project; LHG, TM and AE wrote the manuscript. All authors revised the manuscript.

## Supplemental Methods

### Bioinformatic analysis

For ‘The Cancer Genome Atlas’ (TCGA) dataset Agilent-4502A microarray, data of 488 glioblastoma patients and associated clinical data were downloaded from GlioVis data portal (https://gliovis.bioinfo.cnio.es) (1). Cohort was split into 2 groups of patients defined by the level of Slit2 expression. Overall survival (in months) was used to estimate survival distributions using the Kaplan–Meier method and the distributions were compared using the log-rank test.

### Patient Samples

For the patient samples analyzed in Figure 1C/D and Supplemental Figure 1 I-N, central review histopathology of the patients classified the samples as follows:

45 patients were diagnosed with glioblastoma multiforme (GBM) grade IV, 18 patients with primary anaplastic oligodendroglioma grade III, 6 patients with primary anaplastic astrocytoma grade III, 1 patient with primary anaplastic oligoastrocytoma grade III, 16 patients with grade III mixed anaplasic gliomas, 26 patients with primary oligodendroglioma grade II, 1 patient with recurrent oligodendroglioma grade II, 4 patients with grade II astrocytomas, 9 patients with grade II mixed gliomas, 1 patient with primary xanthoastrocytoma grade II and 1 patient with primary subependymoma grade I. Associated IDH-1/2 mutation status and relevant clinical data from all the 129 patients were used in this study.

### Patient-derived GBM xenograft model (PDX)

N15-0460 patient-derived cell line (PDCL) was established by Gliotex team from GBM tissue sample that was provided by the neuropathology laboratory of Pitie-Salpetriere University Hospital, and obtained as part of routine resections from patients under their informed consent (ethical approval number AC-2013-1962). The parental tumor was IDH-WT and MGMT methylated. Cells are cultivated in DMEM/F12 supplemented with B27, EGF (20 ng/ml), FGF (20 ng/ml), penicillin/streptomycin 1% and plasmocin 0.2%, and dissociated with Accutase. Cells were transduced with luciferase/mkate2 lentiviral particles (in-house produced) at MOI of 3 then shRNA-GFP lentiviral particles (SLIT2, Locus ID 9353, Origene TL309262V) at MOI of 3. After infection, cells were polyclonally selected by Puromycin and mKate^+^ and GFP^+^ cells were sorted by FACS (BioRad S3e Cell Sorter).

For intracranial xenografts, 1.4 x 10^5^ cells were injected in 2 µL of HBSS in Hsd:Athymic Nude-Foxn1nu mice (Envigo) by stereotaxic injection at Bregma AP : +0.1 ; ML : -0.15 ; DV : -0.25 under isoflurane anesthesia (protocol #17503 2018111214011311 v5). Tumor growth was monitored every 15 days by bioluminescence imaging following 100µL luciferin subcutaneous injection at 30mg/mL, and image acquisition with IVIS® Spectrum *in vivo* imaging system (Perkin Elmer). The development of tumors (Tumor take) was evaluated by determining the day when bioluminescence signal doubled compared to the first bioluminescence measured 8 days post-graft.

### *In vitro* spheroid formation and invasion assays

For spheroid formation, 1,000 N15-0460 shCTRL or shSLIT2 cells were plated in non-adherent 96 well plates for 48hs and then imaged by fluorescence using a standard FITC filter to detect endogenous GFP. For invasion assays, spheroids were then resuspended in fibrinogen solution (2.5 mg/ml fibrinogen (Sigma) in DMEM/F12 supplemented with B27, EGF (20 ng/ml), FGF (20 ng/ml) and 50 mg/ml aprotinin (Sigma)) and clotted with 1 U thrombin (Sigma-Aldrich) for 20 min at 37 °C. Cultures were topped with medium and incubated at 37 °C, 5% CO2. After 1 and 2 days, cultures were imaged by fluorescence using a standard FITC filter.

### Extraction of tumor-associated macrophages, lymphocytes and endothelial cells and qPCR analysis

Ketamine/Xylazine anaesthetized tumor-bearing mice were transcardially perfused with 30 ml of ice-cold PBS. Tumors were harvested and incubated with DMEM containing 2.5 mg/ml collagenase D, and 5 U/ml DNase I for 20 min at 37°C. The digested tissue was passed through a 40μm nylon cell strainer (Falcon) and red blood cells were lysed (Red Blood Cells Lysis buffer, Merck).

After blocking with mouse FcR Blocking Reagent (MACS Miltenyi Biotec) cells were stained with the following monoclonal antibodies: anti-CD45 BUV 805 (BD), anti-CD11b BV450 (BD), anti-CD31 PE/CF594 (BD), and anti-CD3 BUV395 (BD), anti-CD4 PE (BD) and anti-CD8 PerCP/Cy5.5 (BD) antibodies. TAMs (CD45^+^CD11b^+^CD3^-^), TALs (CD45^+^CD11b^-^CD3^+^, either CD4^+^ or CD8^+^), endothelial cells (CD45^-^CD31^+^) and tumor cells (CD45^-^GFP^+^) were sorted on a BD FACS Aria II. The cells were then shock-frozen in liquid nitrogen and stored at -80°C until further use.

Total RNA was isolated using the NucleoSpin RNA XS kit from Macherey-Nagel. For protein extraction, frozen cells were resuspended in RIPA Buffer with protease and phosphatase inhibitors and sonicated 3x for 15 seconds each time. Protein concentration was determined by the BCA method and ELISAs were performed according to the manufacturer’s instructions (Mouse VEGF, IL-10 and IFNγ DuoSet ELISA, R&D Systems).

### Cell Growth Determination

Cell viability was determined using the Cell Growth Determination Kit, MTT based (Sigma) according to manufacturer’s instructions. Briefly, 20.000 cells were plated in 24 well plates and grown in normal supplemented medium over 3 days, for determination of their growth curve. After each 24-hour period, cells were incubated with 10% MTT solution for 3 hours, then MTT formazan crystals were dissolved and absorbance was spectrophotometrically measure at 570 nm. Background absorbance measured at 690 nm was subtracted to the first value.

For TMZ sensitivity test, cells were treated for 24 hours with increasing concentrations of TMZ in serum-free medium. The same procedure was performed on untreated cells, and values were normalized and expressed in comparison to untreated cells.

### Vessel perfusion and permeability assay

Glioma-bearing mice from 3 weeks growth were anesthetized and injected intravenously with 100 μL of Alexa Fluor 647 labeled 2,000,000 MW dextran (Life Technologies). Blood vessel perfusion was visualized in vivo using the live imaging settings.

For Miles assay, glioma-bearing mice were anesthetized and injected intravenously with 100 μL 1% Evan’s blue solution (Sigma). Thirty minutes after injection, mice were sacrificed and transcardially perfused with 2% PFA solution. Dissected tumors were weighed and incubated in formamide solution at 56°C overnight to extract the dye. The absorbance of the solution was measured with a spectrophotometer at 620 nm. Five mice per group were analysed. Data are expressed as fold change compared to shCTRL glioma growth with tumor weight normalization.

### Immunofluorescence staining

Vibratome sections of tumors injected in ROSA^mTmG^ reporter mice were blocked and permeabilized in TNBT buffer (0.1 M Tris pH 7.4; NaCl 150 mM; 0.5% blocking reagent from Perkin Elmer, 0.5% Triton X-100) overnight at 4°C. Tissues were then incubated with primary antibodies anti-F4/80 (Life Technologies, 1:100), anti-MRC1 (R&D Systems, 1:100), anti-CD3 (R&D Systems, 1:100), anti-MHCII (Thermo Scientific, 1:100), anti-Glut1 (Millipore, 1:200), anti-Iba1 (Wako, 1:200), anti-Ki67 (Abcam, 1:200), anti-pH2AX (Cell Signaling, 1:100) diluted in TNBT overnight at 4°C, washed in TNT buffer (0.1 M Tris pH 7.4; NaCl 150 mM; 0.5% Triton X-100) at least 7 times and incubated with appropriate Alexa Fluor 647 conjugated antibody (Life Technologies, 1:400) diluted in TNBT overnight at 4°C. Samples were then washed at least 7 times in TNT and mounted on slices in fluorescent mounting medium (Dako). Images were acquired using a Leica SP8 inverted confocal microscope.

### Soluble Flt-1 binding assay

For detection of VEGF expression, vibratome sections were blocked and permeabilized in TNBT overnight at 4°C. Tissues were then incubated with 1μg/ml recombinant mouse soluble Flt-1 FC chimera (R&D Systems) diluted in TNBT for 2.5 h at room temperature. Samples were rinsed three times in TNT and subjected to 4% PFA fixation for 3 min. Samples were washed at least 7 times in TNT and incubated in Alexa Fluor 647 coupled anti-human IgG secondary antibodies (Life Technologies, 1:200) diluted in TNBT overnight at 4°C. Tissues were washed at least 7 times and mounted on slides in fluorescent mounting medium (Dako). Images were acquired using a Leica SP8 inverted confocal microscope.

### Flow-cytometric analysis of tumor-antigen in lymph node immune cells

Deep cervical and mandibular lymph nodes (DCLN and MLN) were dissected from tumor bearing mice 21 days after injection of CT-2A BFP or CT-2A GFP tumor spheroids. The 2 DCLNs and 6 MLNs of each mice were pooled for analysis. LNs were digested for 30 minutes in 1mg/mL Collagenase I diluted in DMEM at 37°C and after RBC lysis, single cell suspensions were prepared by filtering dissociated tissue on 40uM nylon cell strainers. Single cell suspensions were incubated with anti-CD45 APC or BUV805 (BD), anti-CD11b BV650 or BV450 (BD) antibodies. As a control, cells were stained with the appropriate isotype control. Data acquisition was performed on the BD LSRFortessa X20 and analysis was performed with FlowJo_V10.

### Primary cell cultures

Bone-marrow derived macrophages (BMDMs) were isolated from C57BL/6 mice by flushing the femur and tibia with PBS. The bone marrow cells were resuspended in DMEM GlutaMax (Gibco) containing 1% Pen/Strep (Gibco), 20% FBS (Gibco) and 100 ng/mL M-CSF (R&D Systems). Cells were incubated for 2 days at 37 °C and 5% CO2 in non-treated bacterial dishes for adhesion of bone-marrow resident macrophages, and then changed for treated plastic dishes and culture for 6 days with medium change every 2 days. Before experiments, cells were starved in serum- and CSF-free medium overnight. For PI3Kγ inhibition experiments, cells were pre-treated with 1uM IPI-549 for 30 minutes as previously described (2) and then treatments were performed as described for all other experiments.

Microglial cells were obtained as described previously (3, 4). Peritoneal macrophages (PMs) were isolated from peritoneal lavage as previously described (5).

### Immunoprecipitation

After Slit2 treatments for 15 minutes, BMDMs were lysed using NP40 lysis buffer (Boston bioproducts, BP-119X) supplemented with protease and phosphatase inhibitor cocktails (Roche, 11836170001 and 4906845001). Protein concentrations were quantified by BCA assay (Thermo Scientific, 23225) according to the manufacturer’s instructions. 300ug of protein were diluted in 1ml of NP40 buffer containing protease and phosphatase inhibitors for each condition. In the meantime, protein A/G magnetic beads (Thermo fischer, 88802) were washed 5x 10min with NP40 buffer. Protein lysates were incubated for 2 hours at 4°C under gentle rotation with 10ug of PI3Kγ antibodies (Cell Signaling Tecnologies). Then, 50ul of A/G magnetic beads were added to each protein lysate for 2 hours at 4°C under gentle rotation. Beads were then isolated using magnetic separator (Invitrogen) and washed 5 x with NP40 buffer. After the last wash, supernatants were removed and beads were resuspended in 40ul of Laemmli buffer (Bio-Rad, 1610747), boiled at 95°C for 5min and loaded onto 4-15% gradient gels. Western blotting was performed as described above.

### GFP^+^ macrophage isolation

We collected mouse femoral bone-marrows (BMs) before the sacrifice of tumor-bearing mice as previously described for BMDM cultures. In the meantime, rabbit anti-GFP antibodies (Invitrogen) were incubated with sheep anti-rabbit IgG magnetic dynabeads (Invitrogen) in a solution of sterile PBS 0.1%BSA (120ul of beads, 24ul of antibodies in 12ml PBS 0.1%BSA). Solutions were place under gentle rotation at room temperature for 2hours to allow proper coupling of antibodies and beads. Coupled beads were next isolated using a magnetic separator and incubated in the resuspended BMs for 30min. After 5 washes with PBS 0.1%BSA, beads were separated using magnetic separator and RNA was extracted as previously described using RNeasy-kit (Qiagen). RNA samples were and reverse transcribed using SuperScript IV RT (Invitrogen) for gene-deletion verification by qPCR.

## Supplemental Figures

**Supplemental Figure 1.**
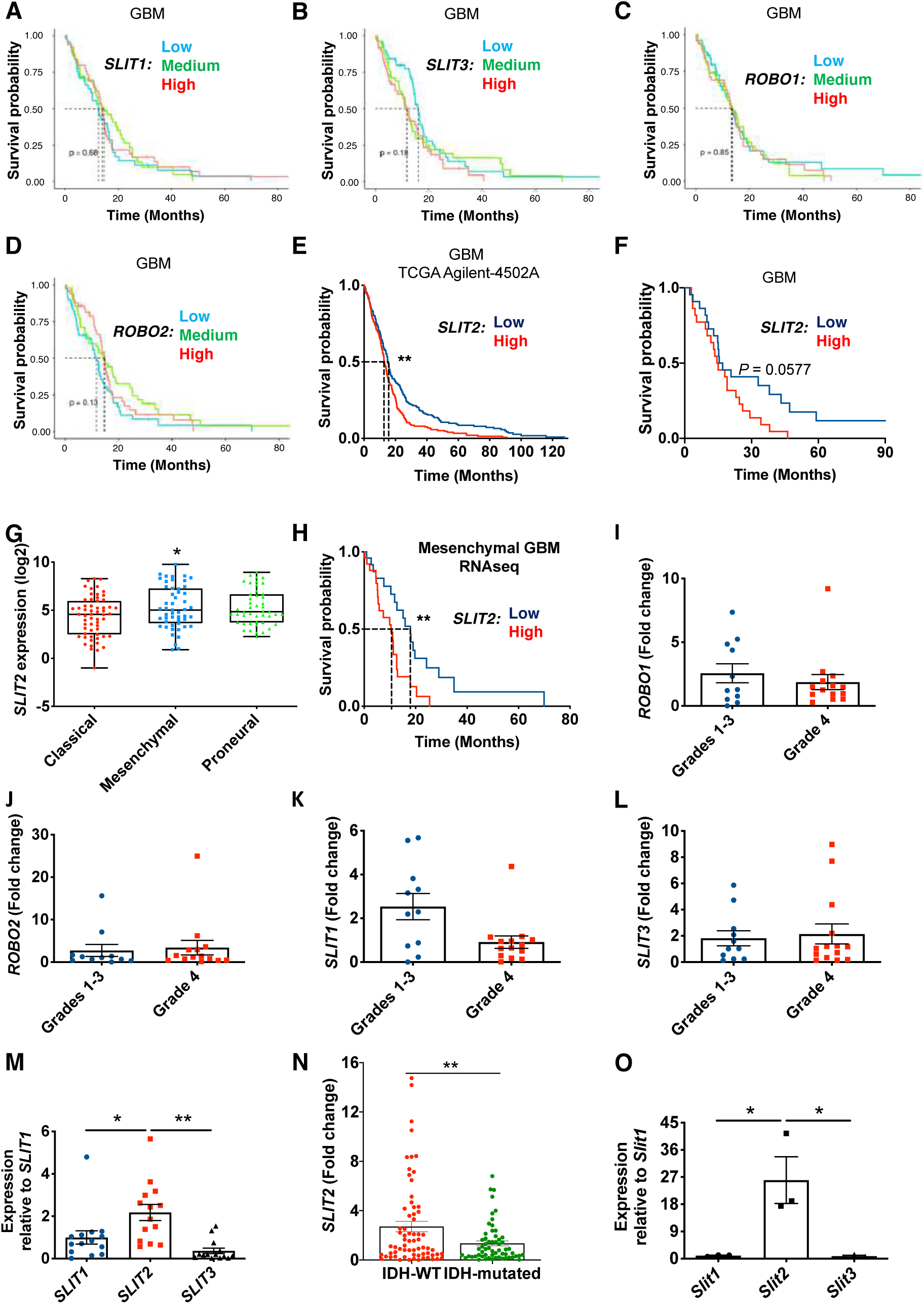
Impact of Slit2 on GBM patient survival. **A-D.** *In silico* analysis of TCGA glioblastoma RNAseq patient database demonstrating that expression of *SLIT1* (**A**), *SLIT3* (**B**), *ROBO1* (**C**) or *ROBO2* (**D**) do not affect patient survival. **E.** *In silico* analysis of TCGA GBM patient microarray Agilent-4502A database showing expression of *SLIT2* is significantly associated with decreased patient survival (*n* = 244 high and 244 low *SLIT2* expressing patients; O.S., 12.9 months for high expression and 15.1 months for low expression, log-rank test). **F.** Survival analysis of GBM patients from (**Figure 1C**) grouped by their levels of *SLIT2* expression (*n =* 22 high and 22 low Slit2 expressing patients; O.S., 14.75 months for high expression and 16.25 months for low expression, log-rank test). **G.** *In silico* analysis of TCGA glioblastoma RNAseq patient database demonstrating *SLIT2* expression in different GBM molecular subtypes (*n =* 59 classical, 51 mesenchymal and 46 proneural tumors; One-Way ANOVA). **H.** *In silico* analysis of TCGA glioblastoma RNAseq patient database demonstrating that *SLIT2* expression is significantly associated with decreased patient survival in mesenchymal GBM patients (*n =* 26 high and 25 low *SLIT2* expressing patients; O.S., 10.4 months for high expression and 17.9 months for low expression, log-rank test). **I-L.** qPCR expression of *ROBO1* (**I)**, *ROBO2* (**J**), *SLIT1* (**K**) and *SLIT3* (**L**) in glioma patient samples (GBM, *n =* 45; LGG, *n =* 84; Student’s t test). **M**. qPCR comparison of *SLIT1*, *SLIT2* and *SLIT3* expression in GBM patient samples (Grade IV, *n =* 14 patients; One-Way ANOVA). **N.** *SLIT2* qPCR expression in all glioma patient samples from (**Figure 1C**) classified by their IDH-1/2 status (IDH-WT, *n =* 67; IDH-mutated, *n =* 59; Student’s t test). **O.** qPCR comparison of *Slit1, Slit2* and *Slit3* expression in CT-2A tumors (*n =* 3 independent tumors, One-Way ANOVA). Data are presented as mean ± s.e.m. * *P* < 0.05, ** *P* < 0.01, *** *P* < 0.001.

**Supplemental Figure 2.**
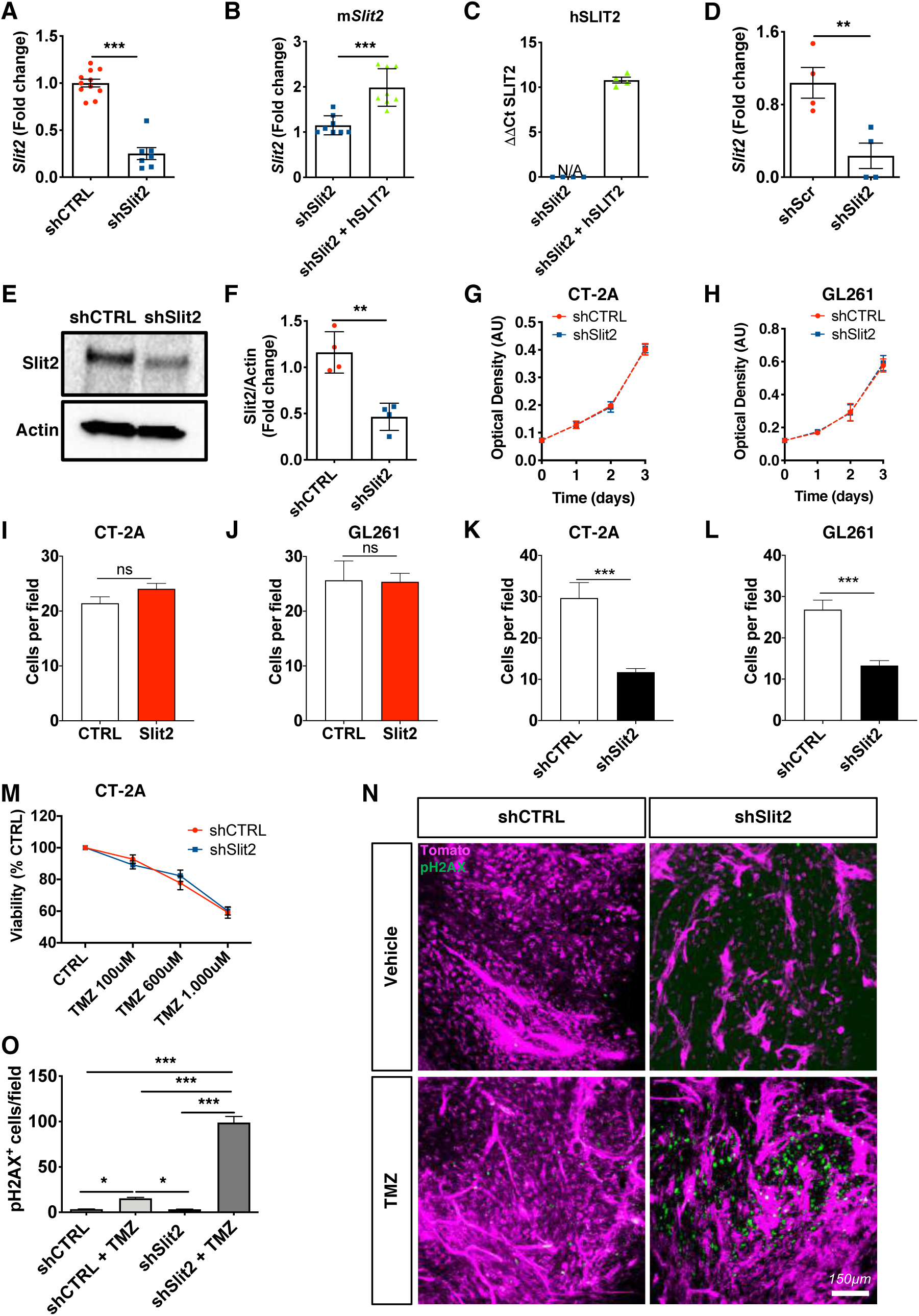
Slit2 silencing does not change tumor cell proliferation or sensitivity to TMZ *in vitro*, but increases TMZ-induced tumor cell death *in vivo*. **A.** *Slit2* qPCR expression in CT-2A shSlit2 and shCTRL (*n =* 10 shCTRL and *n =* 7 shSlit, Student’s t-test). **B-C.** qPCR analysis of murine (**B**, *n =* 8) and human (**C**, *n =* 4) Slit2 expression in cells infected with a human SLIT2 construct (Mann-Whitney U test). **D-F.** qPCR analysis **(D),** western blot analysis **(E)** and protein quantification **(F)** of shRNA Slit2 silencing in GL261 cells (*n =* 4, Mann-Whitney U test). **G-H.** Kinetics of shCTRL and shSlit2 treated CT-2A **(G)** and GL261 **(H)** glioma cell growth over 72 hours in complete medium (*n =* 3, multiple comparison linear regression). **I-J.** Transwell assay quantification of CT-2A (**I**) and GL261 (**J**) cells migration towards a Slit2 gradient (*n =* 4, Mann-Whitney U test). **K-L.** Transwell assay quantification of CT-2A (**K**) and GL261 (**L**) shCTRL or shSlit2 cells invasion towards a serum gradient (*n =* 4, Mann-Whitney U test). **M.** *In vitro* shCTRL and shSlit2 treated CT-2A glioma cell response to TMZ treatment (*n =* 4, One-way ANOVA). **N.** Phospho-H2AX (pH2AX) immunostainings (green) on 23 days tumor sections of CT-2A shCTRL and shSlit2 mice treated or not with TMZ in order to evaluate double-stranded DNA breaks (pH2AX^+^, green) in response to TMZ treatment. **O.** Quantification of **(N)** (*n =* 4 mice per group, 5 fields per tumor, One-way ANOVA). Data are presented as mean ± s.e.m. * *P* < 0.05, ** *P* < 0.01, *** *P* < 0.001.

**Supplemental Figure 3.**
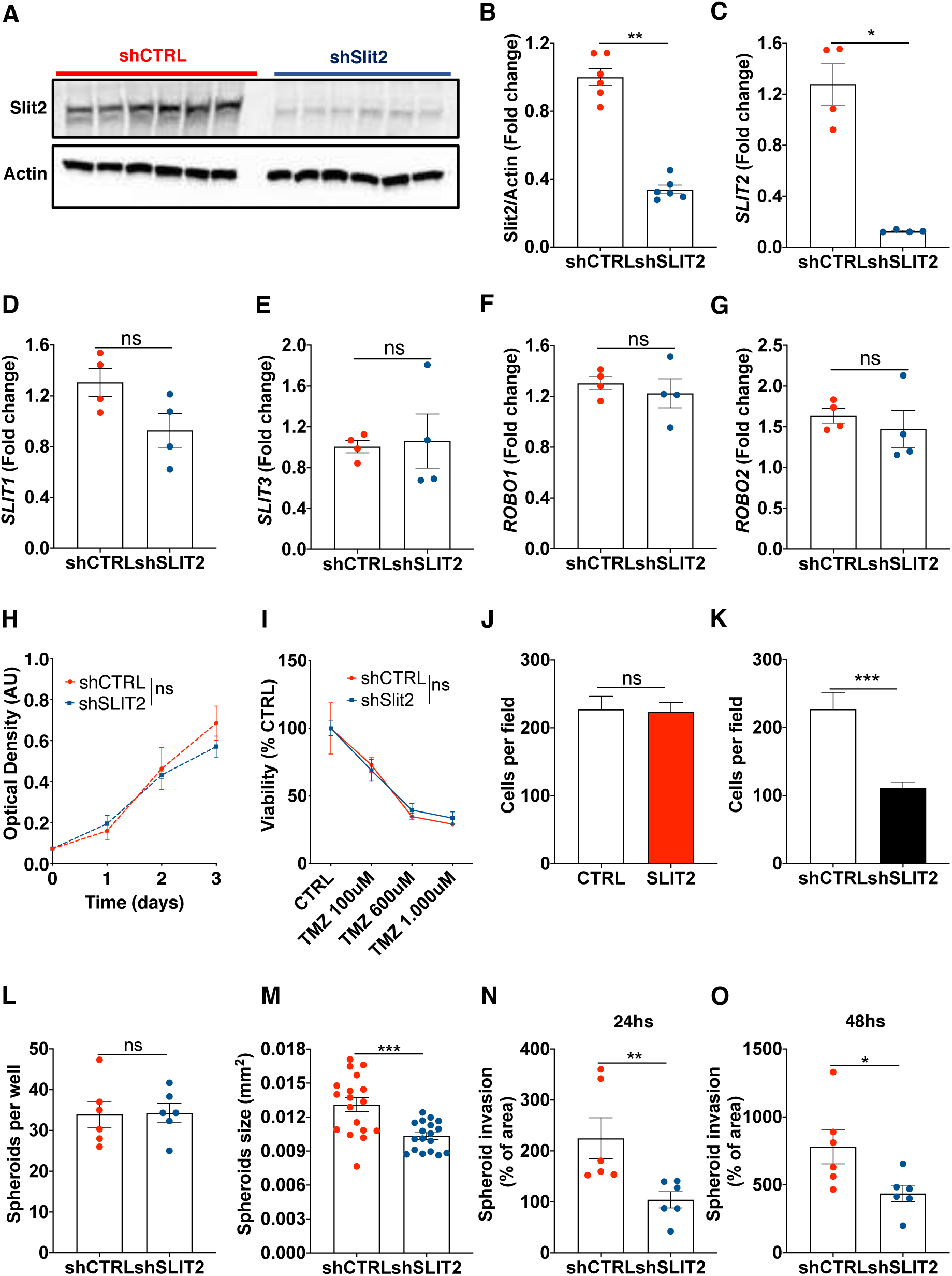
Slit2 silencing reduces invasion of Patient-derived GBM cells. **A-C.** Western blot analysis **(A)**, protein quantification **(B)** and qPCR analysis **(C)** of shRNA *SLIT2* silencing in N15-0460 GBM patient-derived cells (*n =* 6, Mann-Whitney U test). **D-G.** qPCR expression of *SLIT1* (**D**), *SLIT3* (**E**), *ROBO1* (**F**) and *ROBO2* (**G**) in N15-0460 cells after shRNA *SLIT2* silencing (*n =* 4, Mann-Whitney U test). **H.** shCTRL and shSLIT2 treated N15-0460 growth over 72 hours in complete medium (*n =* 3, multiple comparison linear regression). **I.** shCTRL and shSLIT2 treated N15-0460 cells response to TMZ treatment (*n =* 4, Two-way ANOVA). **J.** Transwell assay quantification of N15-0460 cell migration towards a SLIT2 gradient (*n =* 4, Mann-Whitney U test). **K.** Transwell assay quantification of N15-0460 shCTRL or shSLIT2 cell migration towards a serum gradient (*n =* 4, Mann-Whitney U test). **L-M.** Spheroid formation assay quantification of shCTRL and shSLIT2 N15-0460 cells. Number (**L**) and size (**M**) of spheroids formed after 48 hours in culture were quantified (*n =* 6 cultures per group, Mann-Whitney U test). **N-O.** Quantification of spheroid invasion assay in fibrin gels of shCTRL and shSLIT2 N15-0460 cells after 24 (**N**) and 48 hours (**O**). Data are presented as mean ± s.e.m. * *P* < 0.05, ** *P* < 0.01, *** *P* < 0.001.

**Supplemental Figure 4.**
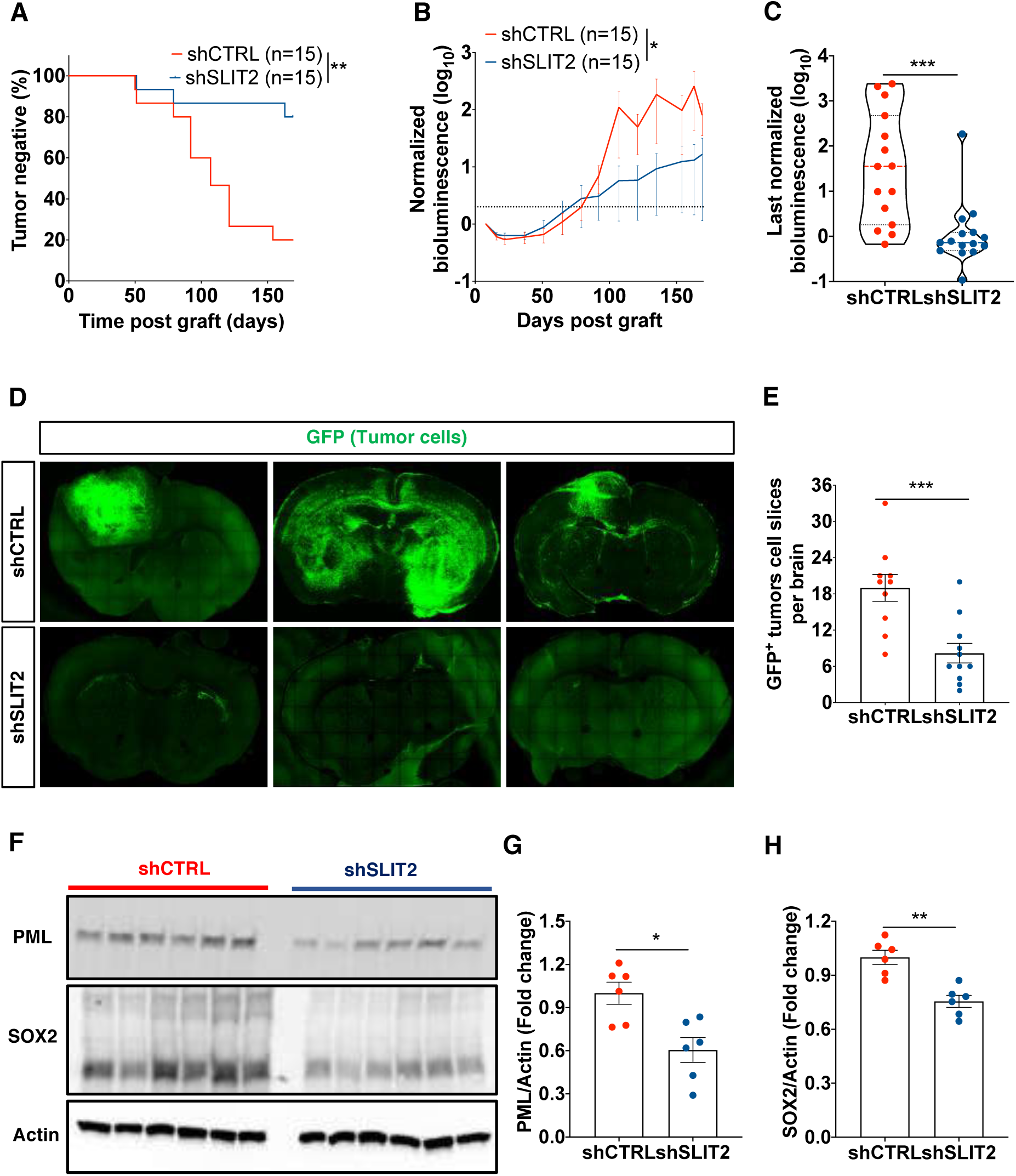
S*L*IT2 silencing slows tumor growth in a GBM Patient-derived Xenograft (PDX) model. **A.** Tumor development curve after injection of shCTRL and shSLIT2 N15-0460 GBM patient-derived cells in nude mice (*n =* 15 mice per group, log-rank test). **B.** bioluminescence signal over time after tumor injection (*n =* 15 mice per group, One-way ANOVA). **C.** bioluminescence signal at the end-point of experiment for each of the injected mice (*n =* 15 mice per group, Mann-Whitney U test). **D.** Tile-scan images of vibratome sections from implanted mice demonstrating GFP^+^ tumor cell spread. **E.** Quantification of GFP^+^ tumor cell spread (*n =* 10 shCTRL and 11 shSLIT2 mice, Mann-Whitney U test). **F-H.** Western blot analysis **(F)** and protein quantification of PML (**G**) and SOX2 (**H**) expression in shCTRL and shSLIT2 N15-0460 GBM cells (*n =* 6, Mann-Whitney U test). Data are presented as mean ± s.e.m. * *P* < 0.05, ** *P* < 0.01, *** *P* < 0.001.

**Supplemental Figure 5.**
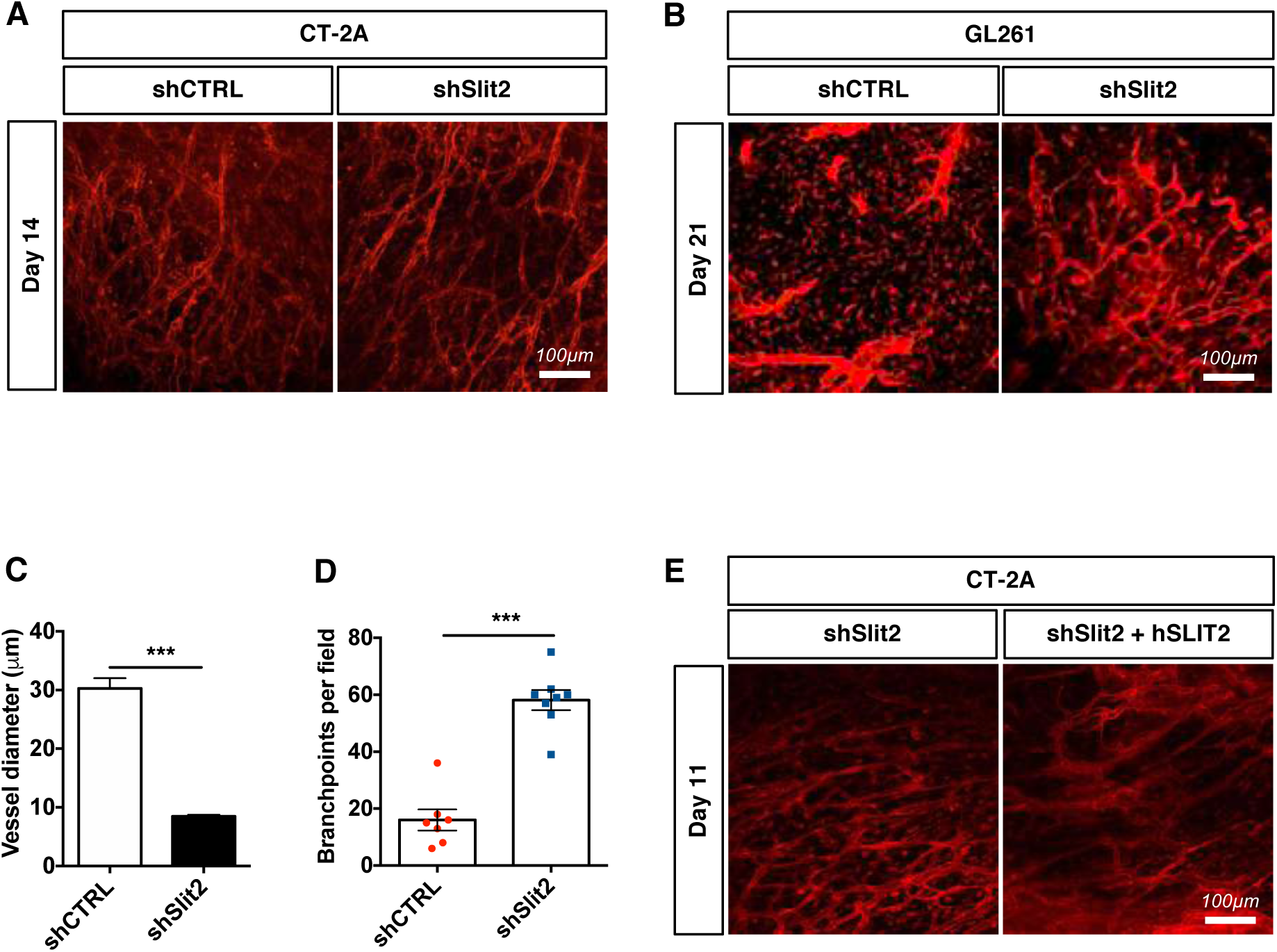
Slit2 drives vessel dysmorphia and vascular dysfunction in CT-2A and GL261 glioma models. **A.** *In vivo* two-photon imaging of ROSA^mTmG^ mice bearing day 14 CT-2A shCTRL or shSlit2 tumors. **B.** *In vivo* two-photon imaging of ROSA^mTmG^ mice bearing day 21 GL261 shCTRL or shSlit2 tumors. **C-D.** Quantification of blood vessel diameter (**C**) and branchpoints (**D**) of the GL261 tumors shown in **(B)** (*n =* 7 mice per group, Student’s t-test). **E.** *In vivo* two-photon imaging of ROSA^mTmG^ mice bearing day 11 CT-2A shSlit2 or shSlit2+hSLIT2 tumors. Data are presented as mean ± s.e.m. * *P* < 0.05, ** *P* < 0.01, *** *P* < 0.001

**Supplemental Figure 6.**
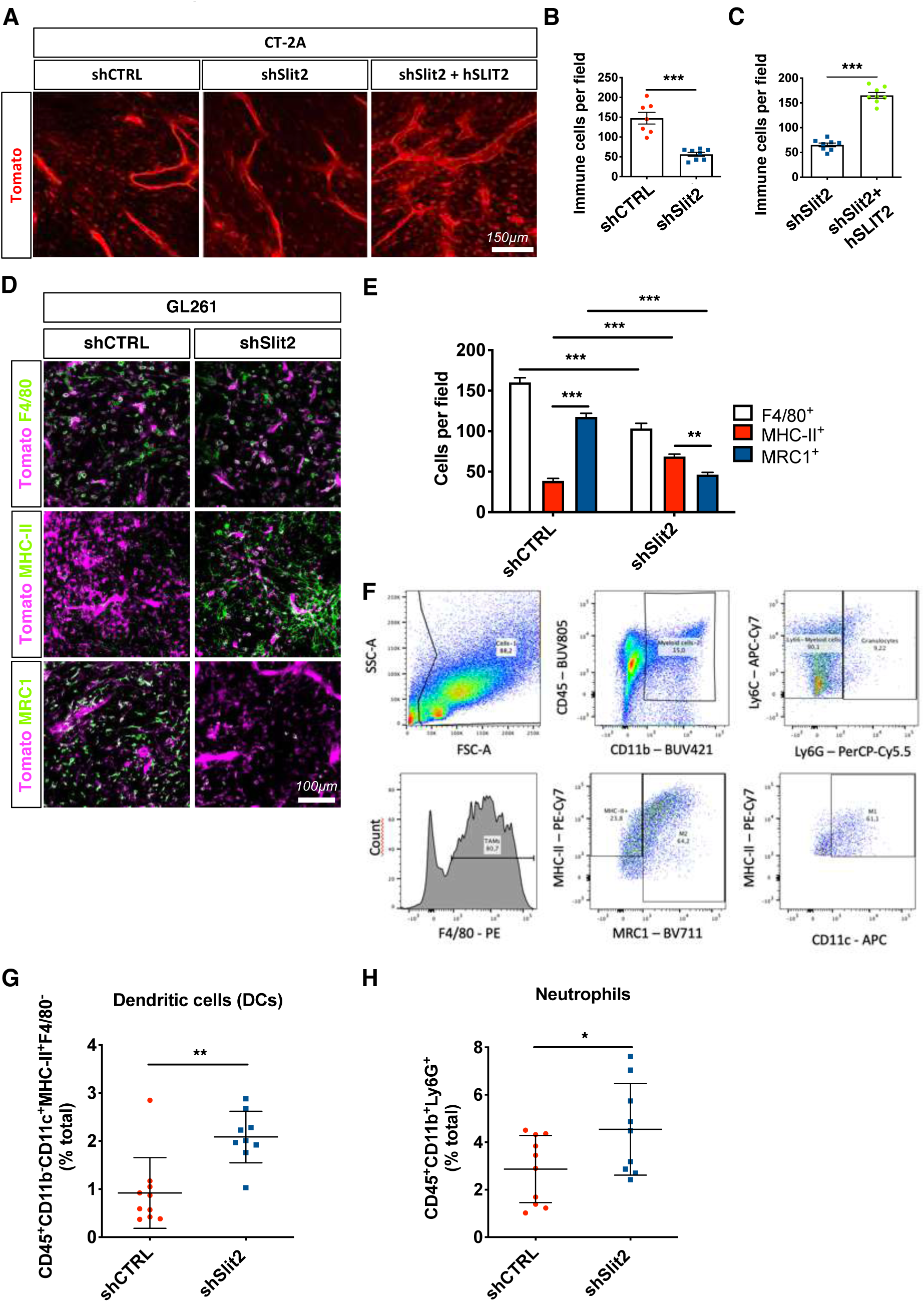
Slit2 silencing favors macrophage cytotoxic polarization in CT-2A and GL261 glioma models. **A-C.** *In vivo* imaging (**A**) and quantification (**B-C**) of host-derived tumor infiltrating immune cells (red) in late-stage CT-2A tumors (*n =* 7 shCTRL, *n =* 8 shSlit2 and shSlit2+hSLIT2 mice, Student’s t-test). **D.** Immunohistochemistry on sections of day 21 GL261 shCTRL or shSlit2 tumors with antibodies recognizing F4/80, MHC-II and MRC1 (green). **E.** Quantifications of (**D)** (*n =* 7 mice per group, 5 fields per tumor, Two-Way ANOVA). **F**. Flow cytometry-gating strategy example for macrophage counting shown in **Figure 4C-E. G-H.** FACS quantification of Dendritic Cells (**G**, DCs, CD45^+^CD11b^+^CD11c^+^MHC-II^+^F4/80^-^) and Neutrophils (**H**, CD45^+^CD11b^+^Ly6G^+^). Data are presented as mean ± s.e.m. * *P* < 0.05, ** *P* < 0.01, *** *P* < 0.001.

**Supplemental Figure 7.**
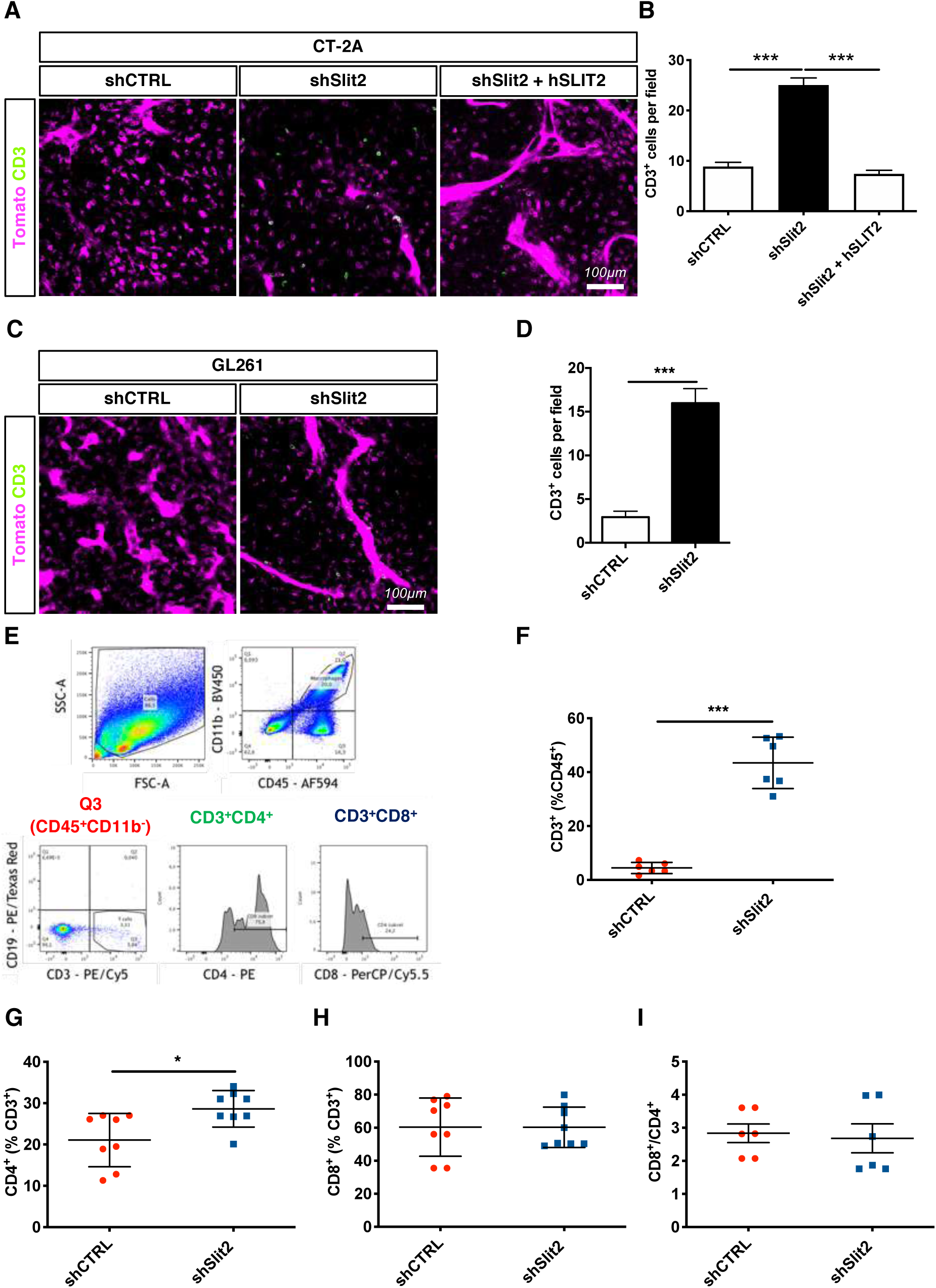
Slit2 drives T cell depletion in CT-2A and GL261 models. **A.** Anti-CD3 staining (green) on sections of late stage CT-2A shCTRL, shSlit2 or shSlit2+hSLIT2 tumors. **B**. Quantification of (**A**) (*n =* 7 mice per group, 5 fields per tumor, One-Way ANOVA). **C**. Anti-CD3 staining (green) on sections of late stage GL261 shCTRL and shSlit2 tumors. **D**. Quantification of (**C**) (*n =* 7 mice per group, 5 fields per tumor, Student’s t-test). **E-H**. Extension of flow-cytometry analysis from **Figure 5**. When considering only the immune cell compartment of the tumor microenvironment (CD45^+^ cells), there is a 10-fold increase in the proportion of TALs (from 4,4% to 43,5%) in shSlit2 tumors **(F)**. Analysis of the percentage of CD4^+^ T helper cells **(G)** and CD8^+^ cytotoxic T cells **(H)** among the TALs (*n =* 8 mice per group, Mann-Whitney). **I.** Ratio between CD8^+^ and CD4^+^ TALS (*n =* 8 mice per group, Mann-Whitney). Data are presented as mean ± s.e.m. * *P* < 0.05, ** *P* < 0.01, *** *P* < 0.001.

**Supplemental Figure 8.**
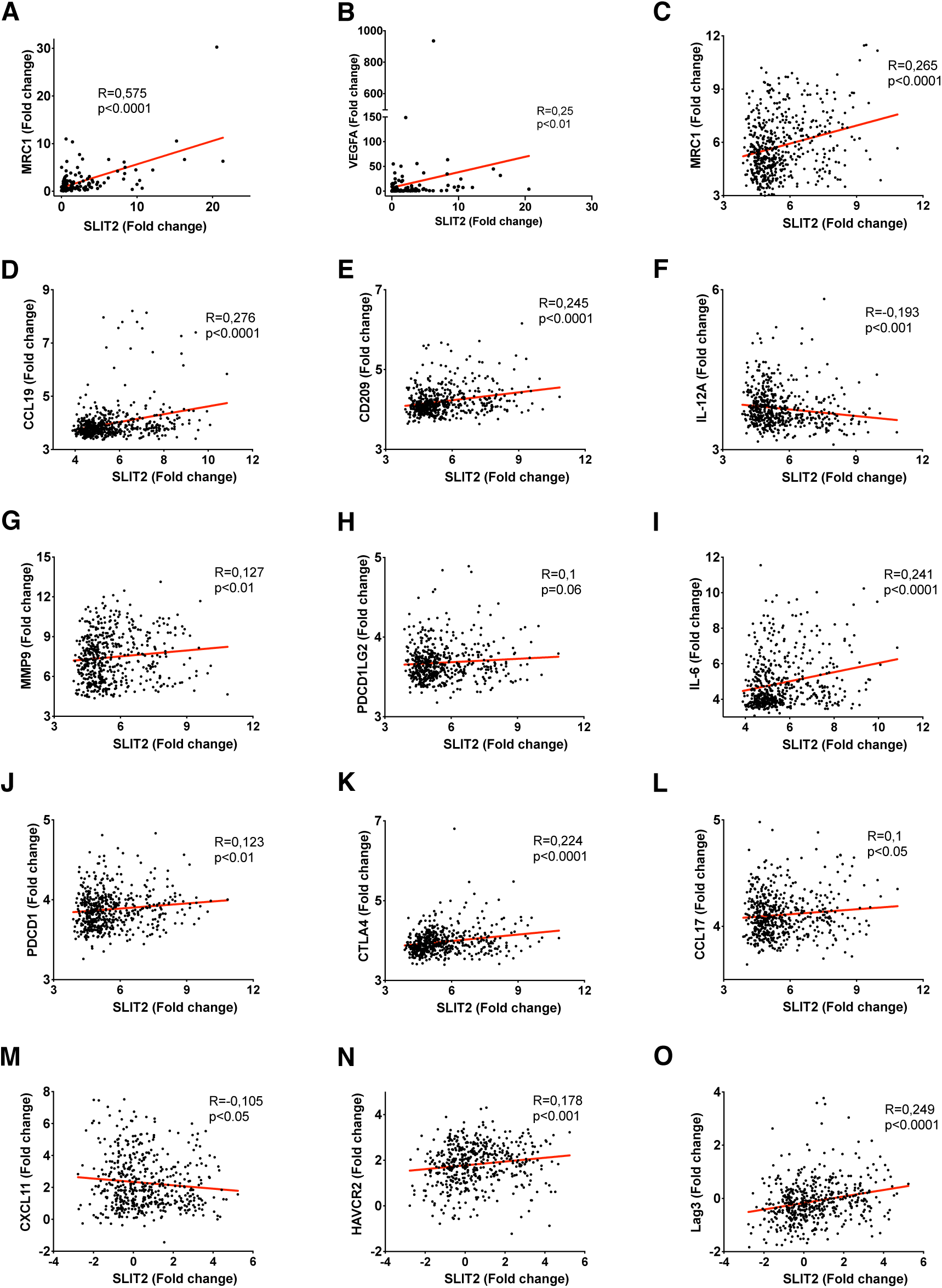
Slit2 expression correlated with immunosuppression in GBM patients. **A-B.** Correlation analysis of MRC1 (**A)** or VEGFA (**B**) and SLIT2 expression in GBM patients (*n =* 129 patients, Spearman’s correlation test). **C-O.** Correlation analysis of SLIT2 expression with the indicated genes in GBM patients from TCGA cohort (*n =* 489 patients, Spearman’s correlation test). * *P* < 0.05, ** *P* < 0.01, *** *P* < 0.001.

**Supplemental Figure 9.**
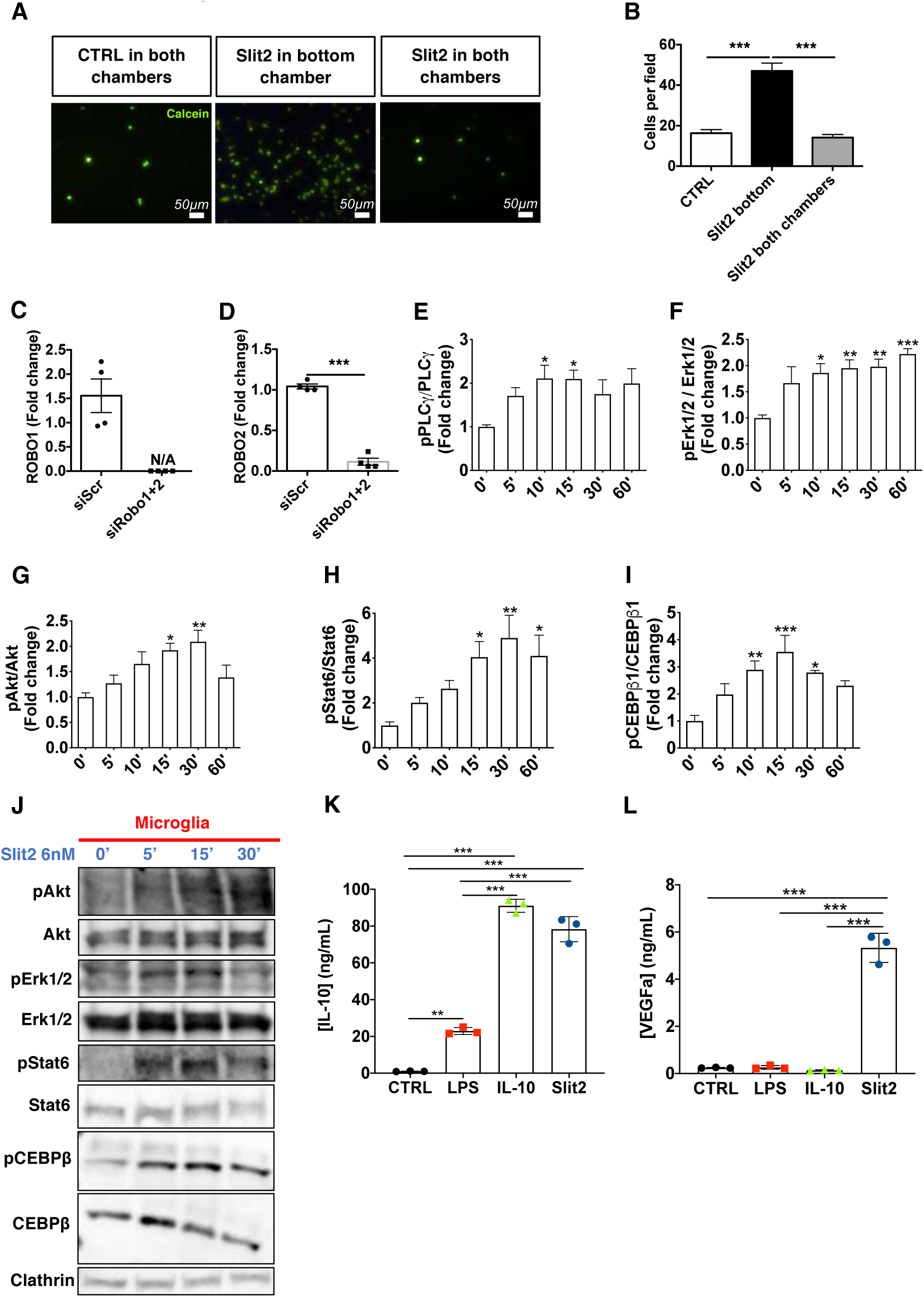
Slit2 induces chemotactic migration and signaling in primary macrophages and microglial cells. **A-B.** Representative images **(A)** of calcein-stained (green) Transwell assays with Slit2 treatment in the bottom chamber or in both chambers and quantification **(B)**. Slit2- induced migration is chemotactic, as treatment with Slit2 in both chambers disrupts the gradient and abrogates migration (*n =* 4, One-way ANOVA). **C-D.** qPCR analysis of Robo1 (**C**) and Robo2 (**D**) expression after siRNA treatment of cultured RAW264.7 macrophages for 72hs (*n =* 4, Mann Whitney). **E-I.** Quantifications of the Western Blots shown in **Figure 6F**. (*n =* 6, One-Way ANOVA). **J**. Western blot analysis of Slit2 downstream signaling in cultured microglial cells (*n =* 3). **K-L.** ELISA from conditioned medium from LPS or Slit2-treated microglial cells quantifying the secretion of IL-10 (**K**) and VEGFa (**L**) (*n =* 3 independent cultures, Mann-Whitney U test). Data are presented as mean ± s.e.m. * *P* < 0.05, ** *P* < 0.01, *** *P* < 0.001.

**Supplemental Figure 10.**
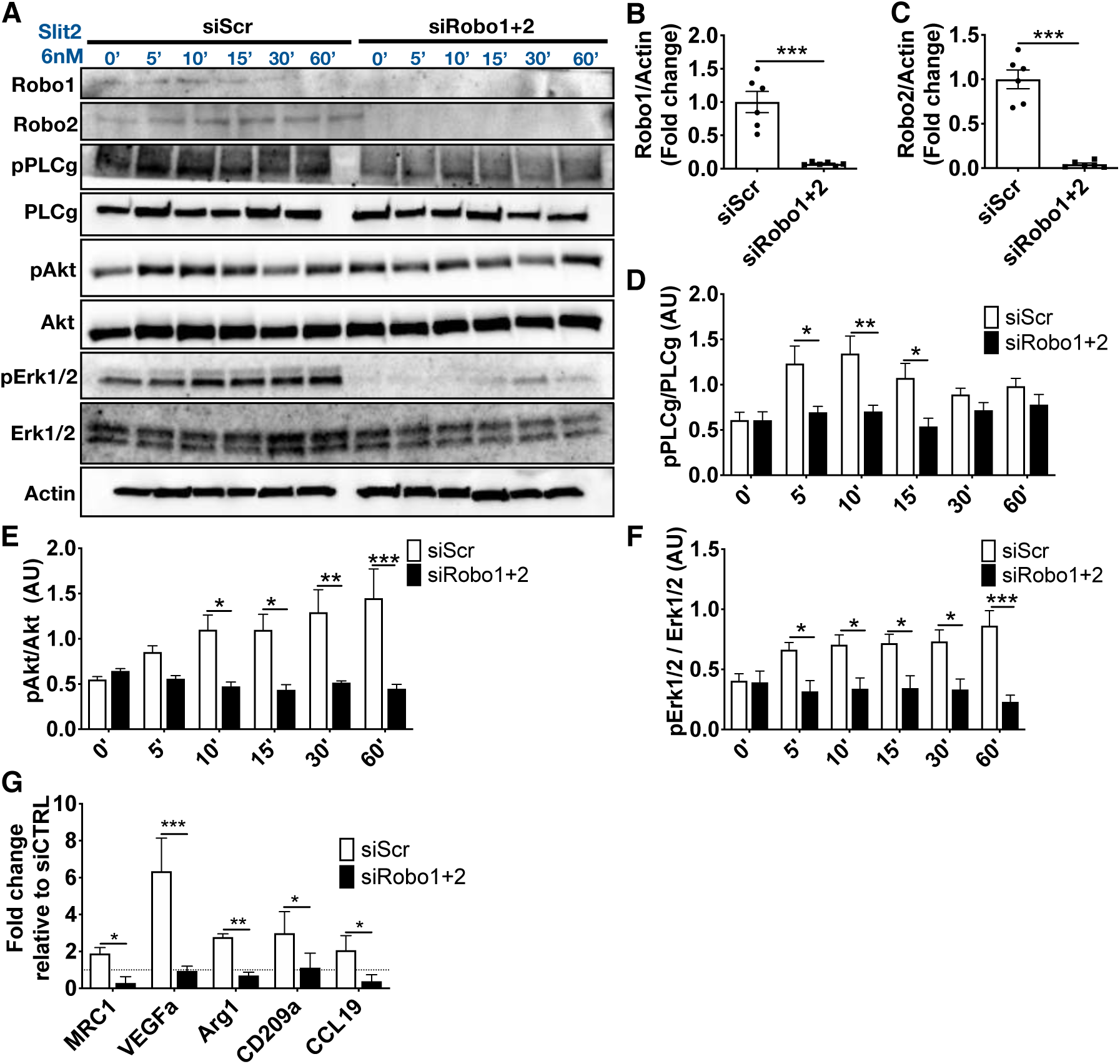
Slit2 induces macrophage migration and downstream signaling via Robo1/2. **A.** Western blot analysis of PLCγ, Akt and Erk1/2 phosphorylation induced by Slit2 in control and Robo1/2 knockdown RAW264.7 macrophages (*n =* 5). **B-C**. Quantification of Robo1 (**A**) and Robo2 (**B**) protein expression after Robo1 and Robo2 knockdown. **D-F**. Quantification of **(A)** (*n =* 5, Two-way ANOVA). **G.** qPCR analysis of genes related to the tumor supportive phenotype (*Mrc1*, *Vegfa, Arg1, Cd209a* and *Ccl19*) in RAW264.7 macrophages after Robo1 and Robo2 knockdown and Slit2 treatment (*n =* 4, Mann Whitney U test). Data are presented as mean ± s.e.m. * *P* < 0.05, ** *P* < 0.01, *** *P* < 0.001.

**Supplemental Figure 11.**
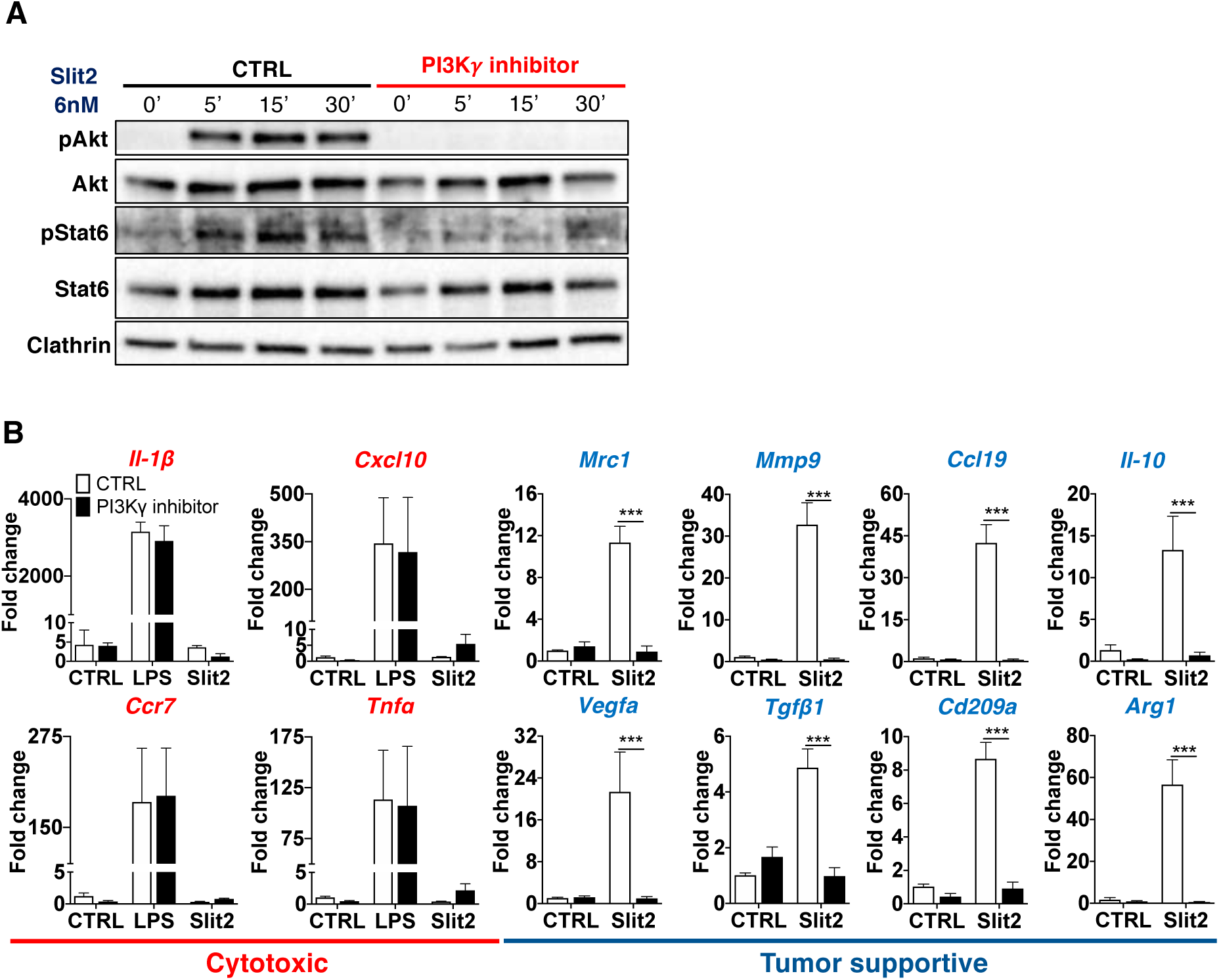
PI3Kγ inhibiton disrupts Slit2-induced macrophage and microglia polarization. **A.** WB analysis of Akt and Stat6 phosphorylation in BMDMs induced by Slit2 after PI3Kγ inhibitor IPI-549 pretreatment (*n =* 3 independent cultures). **B.** qPCR analysis of microglial cultures following Slit2 or LPS treatment after pre-treatment with PI3Kγ inhibitor (*n =* 4 independent cultures, 2-way ANOVA). Data are presented as mean ± s.e.m. * *P* < 0.05, ** *P* < 0.01, *** *P* < 0.001.

**Supplemental Figure 12.**
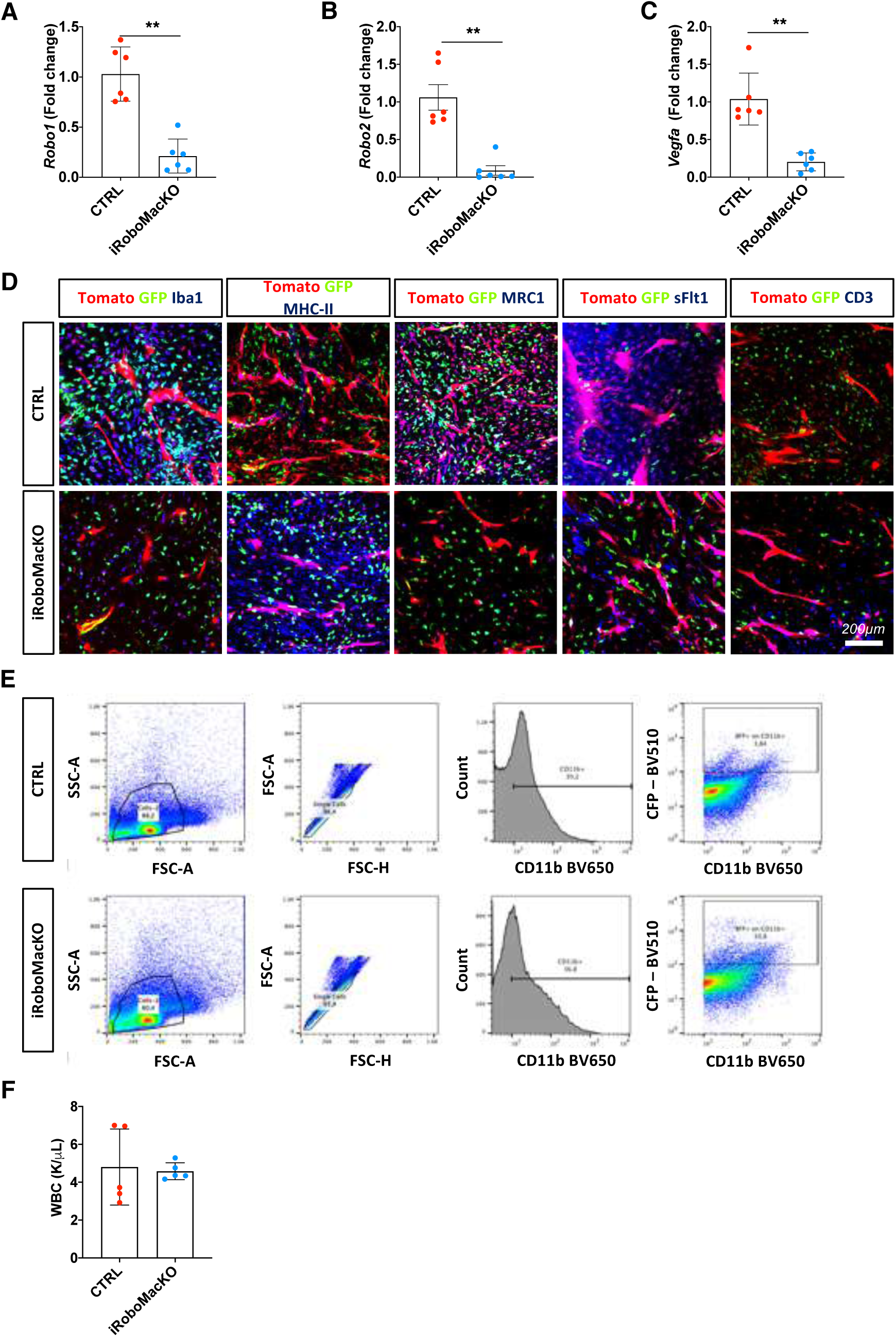
Analysis of iRoboMacKO. **A-C.** qPCR of Robo1 (**A**), Robo2 (**B**) and Vegfa (**C**) in GFP^+^ macrophages extracted from the bone-marrow of CTRL and iRoboMacKO tumor-bearing mice 21 days after tumor implantation. **D**. Immunohistochemistry images related to quantifications shown in **Figure 8I-K**. **E.** Flow cytometry-gating strategy example for graphs shown in **Figure 8M-N**. **F.** Total white blood cells (WBC) counts from peripheral blood of late-stage CTRL and iRoboMacKO tumor-bearing mice (*n =* 5 mice/group; Mann-Whitney U test). Data are presented as mean ± s.e.m. * *P* < 0.05, ** *P* < 0.01, *** *P* < 0.001.

**Supplemental Figure 13.**
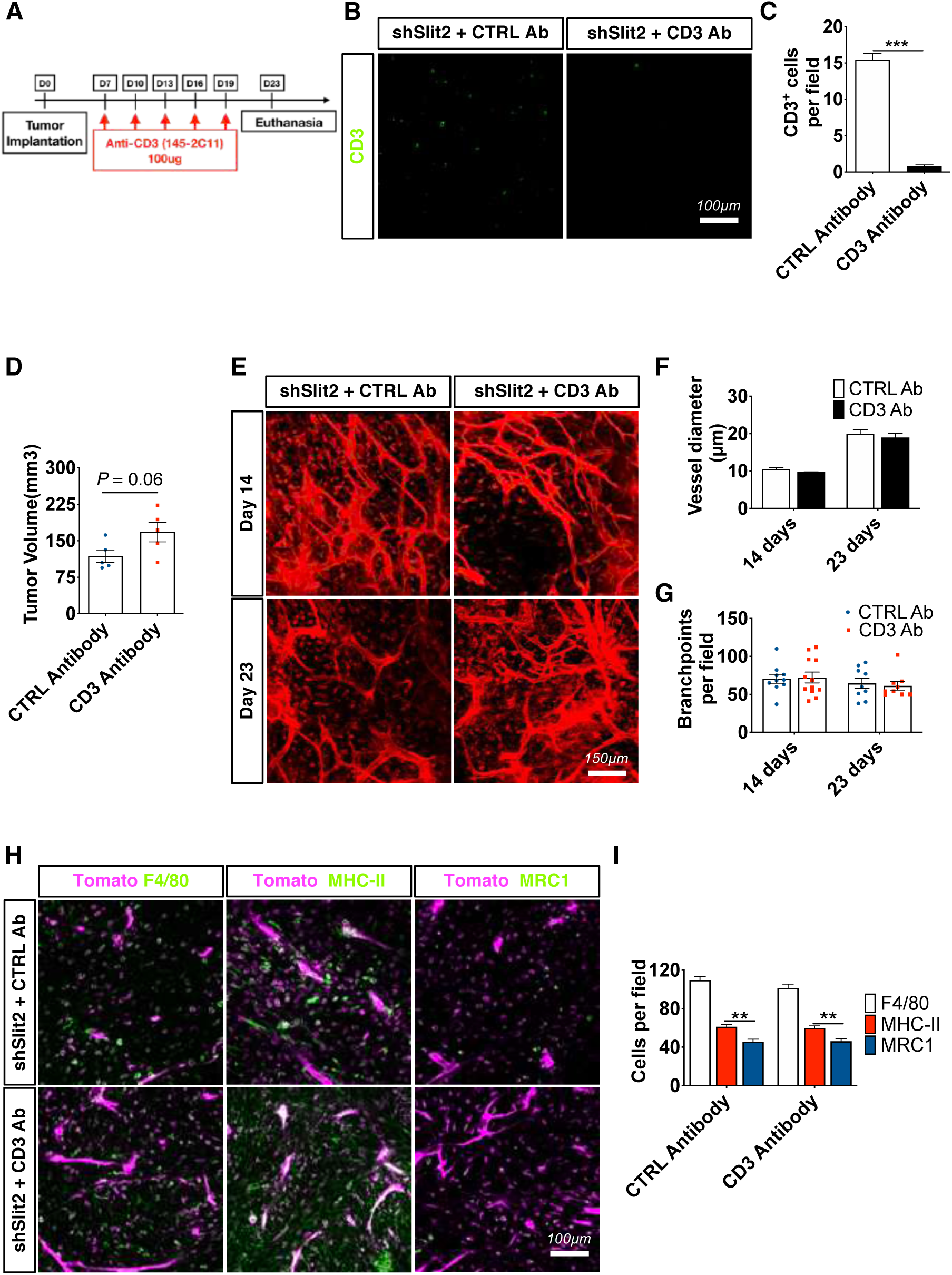
I*n vivo* T cell-depletion does not affect the TME. **A.** Experimental design for T cell depletion by intravenous injection with 145-2C11 anti- CD3 antibodies (mice were treated with 100ug of 145-2C11 antibodies every 3 days starting 7 days after tumor implantation). **B.** CD3 immunostainings performed on sections of late stage CT-2A tumors. **C.** Quantification from **(B)** (*n =* 5 mice per group, 5 fields per staining, Student’s t test). **D.** Tumor volume quantification at 23 days following anti- CD3 mAb treatment (*n =* 5 mice per group, Mann-Whitney U test). **E.** *In vivo* two-photon imaging of ROSA^mTmG^ mice bearing early (14 days) and late stage (23 days) CT-2A shSlit2 tumors with or without anti-CD3 mAb treatment. **F-G.** Quantification of Blood vessel diameter **(F)** and branchpoints **(G)** from **€** (*n =* 5 mice per group, One-way ANOVA). **H.** F4/80, MHC-II and MRC1 Immunohistochemistry on sections of late stage (23 days) CT-2A shSlit2 tumors treated with control mAb or anti-CD3 mAb. **I.** Quantification from (**H**, *n =* 5 mice per group, 5 fields per tumor, Two-way ANOVA). Data are presented as mean ± s.e.m. * *P* < 0.05, ** *P* < 0.01, *** *P* < 0.001.

**Supplemental Figure 14.**
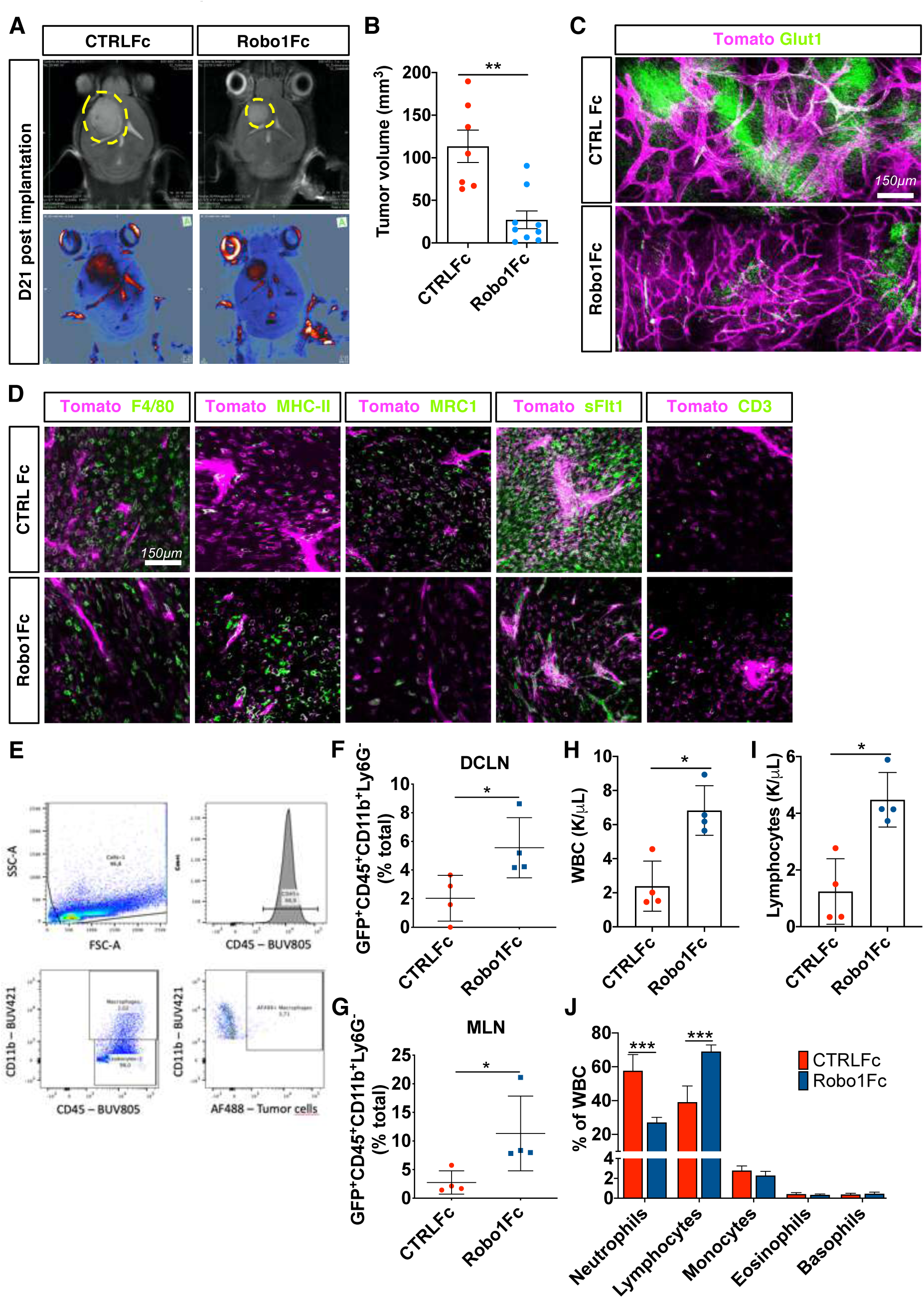
Robo1Fc treatment slowed GBM growth by inducing systemic long-term anti-tumor immune responses. **A.** T2-weighted post-gadolinium MRI images of CTRLFc and Robo1Fc treated mice 21 days after tumor implantation. **B.** Quantification of tumor size from (**A**) (*n =* 7 CTRLFc and 9 Robo1Fc tumors). **C.** Immuno-staining for Glut1 (quantified in **Figure 9H**). **D.** Immunohistochemistry images related to quantifications shown in **Figure 9I-K**. **E**. Flow cytometry-gating strategy example for graphs shown in (**F-G**). **F-G.** FACS analysis of deep cervical and mandibular lymph nodes (DCLN and MLN, respectively) from late-stage CTRLFc- and Robo1Fc-treated mice (*n =* 4 mice/group; Mann-Whitney U test). **H-J.** Total white blood (WBC, **H**), lymphocyte (**I)** and differential WBC (**J**) counts from peripheral blood of late-stage CTRLFc- and Robo1Fc-treated tumor-bearing mice (*n =* 4 mice/group; Mann-Whitney U test and Two-way ANOVA). Data are presented as mean ± s.e.m. * *P* < 0.05, ** *P* < 0.01, *** *P* < 0.001.

**Supplemental Data Table. 1.** List of qPCR Primers used in this study.

| Primer | Cat No |
| --- | --- |
| Hs_ROBO3_1_SG | QT00055951 |
| Hs_ROBO2_2_SG | QT01007664 |
| Hs_ROBO4_1_SG | QT00237741 |
| Hs_SLIT2_1_SG | QT00007784 |
| Hs_SLIT3_1_SG | QT00018795 |
| Hs_SLIT1_1_SG | QT00071113 |
| Hs_ROBO1_2_SG | QT01668982 |
| Hs_ACTB_1_SG | QT00095431 |
| Mm_ACTB_1_SG | QT00095242 |
| Mm_GAPDH_3_SG | QT01658692 |
| Mm_CCR7_1_SG | QT00240975 |
| Mm_MRC1_1_SG | QT00103012 |
| Mm_VEGFA_1_SG | QT00160769 |
| Mm_CCL19_2_SG | QT02532173 |
| Mm_TNF_1_SG | QT00104006 |
| Mm_MMP9_1_SG | QT00108815 |
| Mm_TGFB1_1_SG | QT00145250 |
| Mm_IL1B_2_SG | QT01048355 |
| Mm_PDCD1IG1_1_SG | QT00148617 |
| Mm_PDCD1IG2_1_SG | QT00136640 |
| Mm_CXCL10_1_SG | QT00093436 |
| Mm_IL12B_1_SG | QT00153643 |
| Mm_CD209A_1_SG | QT00116312 |
| Mm_ARG1_1_SG | QT00134288 |
| Mm_IL10_1_SG | QT00106169 |
| Mm_IL12M_1_SG | QT00101108 |
| Mm_IL2_1_SG | QT00112315 |
| Mm_CXCL11_1_SG | QT00265041 |
| Mm_IL17A_1_SG | QT00103278 |
| Mm_IFNg_1_SG | QT01038821 |
| Mm_CCL17_1_SG | QT00131572 |
| Mm_PDCD1_1_SG | QT00111111 |
| Mm_ROBO1_1_SG | QT00146853 |
| Mm_SLIT1_1_SG | QT01044925 |
| Mm_SLIT2_1_SG | QT00163828 |
| Mm_SLIT3_1_SG | QT00283416 |
| Mm_ROBO3_1_SG | QT00136605 |
| Mm_ROBO2_1_SG | QT00143255 |

